# Overcoming artificial structures in resolution-enhanced Hi-C data by signal decomposition and multi-scale attention

**DOI:** 10.1101/2024.10.21.619560

**Authors:** Qinyao Li, Kelly Yichen Li, Chiara Nicoletti, Pier Lorenzo Puri, Qin Cao, Kevin Y. Yip

## Abstract

Computational enhancement is an important strategy for inferring high-resolution features from genome-wide chromosome conformation capture (Hi-C) data, which typically have limited resolution. Deep learning has been highly successful in this task but we show that it creates prevalent artificial structures in the enhanced data due to the need to divide the large contact matrix into small patches. In addition, previous deep learning methods largely focus on local patterns, which cannot fully capture the complexity of Hi-C data. Here we propose Smooth, High-resolution, and Accurate Reconstruction of Patterns (SHARP) for enhancing Hi-C data. It uses the novel approach of decomposing the data into three types of signals, due to one-dimensional proximity, contiguous domains, and other fine structures, and applies deep learning only to the third type of signals, such that enhancement of the first two is unaffected by the patches. For the deep learning part, SHARP uses both local and global attention mechanisms to capture multi-scale contextual information. We compare SHARP with state-of-the-art methods extensively, including application to data from new samples and another species, and show that SHARP has superior performance in terms of resolution enhancement accuracy, avoiding creation of artificial structures, identifying significant interactions, and enrichment in chromatin states.

## Introduction

In mammalian genomes, long DNA molecules fold in the three-dimensional (3D) space into compact structures such that they can be packed into the cell nucleus. Understanding such spatial structures can provide insights into complex relationships between genome architecture, gene activities, and functional states of the cell^1–5^. High-throughput chromosome conformation capture (Hi-C)^6^ is the most commonly used method for probing the overall 3D genome architecture by detecting pair-wise chromatin contacts across the entire genome. The resulting data is represented by a contact matrix, which records interaction frequencies between equally sized genomic bins. Typical bin size used in published studies ranges from 1kb to 1Mb. In order to have sufficiently reliable contact counts, it has been proposed that the resolution of a Hi-C contact matrix is the smallest bin size such that *≥* 80% of the bins have *≥* 1,000 contacts^7^. For the human and mouse genomes, Hi-C data with a resolution of 40kb or lower is usually considered low-resolution^8,9^, whereas data with a resolution of 5kb or higher is usually considered high-resolution^9,10^.

Since a linear increase in the resolution requires a quadratic increase in the number of sequencing reads^11^, few existing Hi-C data sets have very high resolution, but it is required for detecting fine-scaled local structures. High-resolution data are also particularly important for identifying regulatory regions and their contacts with potential target genes in normal and disease states^12–15^ since there can be multiple regulatory elements and genes within the vicinity of a small genomic region.

To reconstruct high-resolution genome structures from low-resolution Hi-C data, computational methods have been employed. The state-of-the-art approach is to treat the Hi-C contact matrix as an image and define the reconstruction problem as image resolution enhancement^8,16–19^, using deep artificial neural network (ANN) models such as convolutional neural networks (CNNs)^20^ or generative adversarial networks (GANs)^21^. Due to the large size of the contact matrix, all these methods have to divide it into small regular-sized submatrices (to be referred to as “patches” hereafter) and apply the model to each of them separately. As to be shown in Results, the boundaries of these patches frequently create discontinuities in the reconstructed high-resolution contact matrix and consequently, artificial structures such as fake topologically associating domains (TADs)^22–24^ that do not actually exist are wrongly called.

In this work, we propose Smooth, High-resolution, and Accurate Reconstruction of Patterns (SHARP), a novel method for enhancing resolution of Hi-C contact matrices that avoids creating artificial structures. We demonstrate that SHARP outperforms current state-of-the-art methods in terms of both resolution enhancement and avoidance of artificial structures across various data sets and settings. As a result, downstream analyses are also more accurate with the SHARP-enhanced Hi-C contact matrices, such as detection of significant interactions and their chromatin states.

## Results

### Overview of SHARP

The artificial structures created by existing Hi-C resolution enhancement methods originate from the fact that the patches they define as input units do not align with the actual structures observed in the contact matrix (Figure 1a). As a result, in the reconstructed highresolution matrix, values on the two sides of a patch boundary can take on very different values since the two patches are enhanced by the model separately (Figure 1b), which is an artifact that does not happen in the actual high-resolution contact matrix (Figure 1c). Some existing methods attempt to solve this problem by defining overlapping patches^18,25^ but that creates substantial computational overhead and adds extra complexity in merging the reconstruction results of overlapping patches. On the other hand, adding a post-hoc smoothing step is also not a good solution since it can destroy real structures that happen to align with the patches.

**Figure 1:**
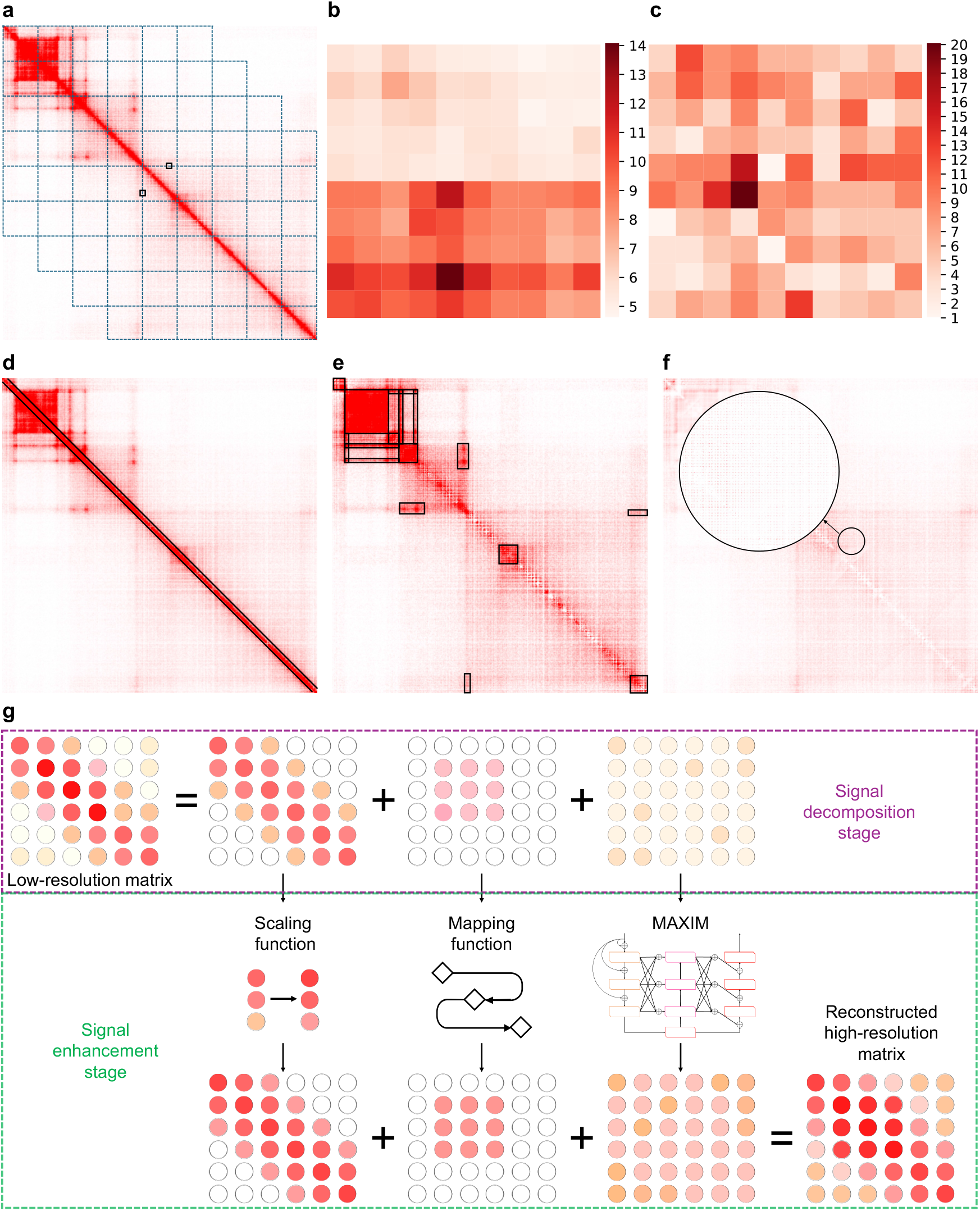
Overview of SHARP. **a**, An example region (Chr20:53.4-56.3Mb) showing that boundaries of 320kb *×* 320kb patches (squares wrapped by blue dashed lines) do not align with real structures in the actual high-resolution GM12878 contact matrix. **b**, Resolution-enhanced contact matrix produced by the published method HiCNN2-3^25^ of an example area in the contact matrix (Chr20:54.935Mb-54.985Mb *×* Chr20:54.695Mb-54.745Mb, marked by the black boxes in Panel a). The horizontal line in the middle is the patch boundary. **c**, The actual high-resolution contact matrix of the same area depicted in Panel b. **d-f**, Decomposition of the contact matrix into signals due to 1D proximity (diagonal band in Panel d), contiguous domains (rectangles in Panel e), and other fine structures (Panel f). The three panels show the original matrix, the matrix after subtracting out signals due to 1D proximity, and the matrix after further subtracting out signals due to contiguous domains, respectively. The large circle in Panel f provides a zoomed-in view of the area indicated by the small circle. **g**, The two stages of SHARP during model training.

SHARP tackles this problem by modeling the Hi-C contact matrix as the superposition of signals caused by three types of chromatin contacts, namely 1) contacts due to one-dimensional (1D) proximity, which create strong signals close to the main diagonal of the matrix^26^ (Figure 1d), 2) contacts due to contiguous domains, which create stronger signals in on-diagonal rectangles (within-domain contacts) and off-diagonal rectangles (between-domain contacts) as compared to their surroundings (Figure 1e), and 3) contacts due to other fine structures such as individual chromatin loops, which create more subtle signals (Figure 1f). We argue that ANN-based resolution enhancement is most useful for the third type of signals due to their intricate properties, while the first two types of signals can be captured by simpler models that do not require defining the patches.

Specifically, the procedure of SHARP is divided into a signal decomposition stage and a signal enhancement stage (Figure 1g, Methods). In the signal decomposition stage, signals due to 1D proximity are modeled by an analytical function^27^ while signals due to contiguous domains are detected by our novel block detection and modeling algorithms (Methods). We allow the blocks to be overlapping and nested due to complex arrangements of structures in the 3D genome. After subtracting out these two types of signals, the remaining contact matrix is expected to be more uniform and contains mostly subtle local structures (Figure 1f). Then in the signal enhancement stage, the three types of signals are enhanced separately and then added together to form the high-resolution matrix: Signals due to 1D proximity are scaled to match the expected level in the high-resolution matrix; signals due to contiguous domains are enhanced by a mapping function; the remaining signals are enhanced using an ANN (Methods). Since the first two types of signals are identified from the whole contact matrix directly without dividing it into patches, the final resolution-enhanced matrix is much less affected by the patch boundaries.

For the ANN, we chose the Multi-Axis Multi-Layer Perceptron for Image Processing (MAXIM) architecture^28^, which uses both local attention and global attention mechanisms to effectively capture multi-scale contextual information and detailed textures, thereby achieving excellent performance in image enhancement tasks. During model training, parameters of the models for all three types of signals are learned based on paired high-resolution and low-resolution contact matrices of the same biological sample. The learned models can then be applied to enhance the resolution of a low-resolution contact matrix from another sample directly.

### SHARP reconstructs high-resolution Hi-C contact matrices accurately

To evaluate the effectiveness of SHARP, following previous studies^8,17–19,25,29–31^, we applied it to enhance a low-resolution contact matrix created by down-sampling an actual high-resolution contact matrix produced from human GM12878 cells^7^. We trained the models using data in both the high-resolution and low-resolution matrices from 15 chromosomes, selected the best model during the training process using 4 other chromosomes, and finally evaluated the enhancement performance using the remaining 3 left-out chromosomes (Chr 3, 10, and 20) (Methods). For benchmarking purpose, we also applied three state-of-the-art Hi-C resolution enhancement methods in the same way, namely HiCARN-2^19^, HiCNN2-3^25^, and HiCPlus^8^. Based on four standard performance measures (Pearson correlation coefficient [PCC], Spearman correlation coefficient [SCC], peak signal-to-noise ratio [PSNR] and structural similarity index measure [SSIM]; Methods), SHARP consistently outperformed these three methods in all cases (Figure 2a).

**Figure 2:**
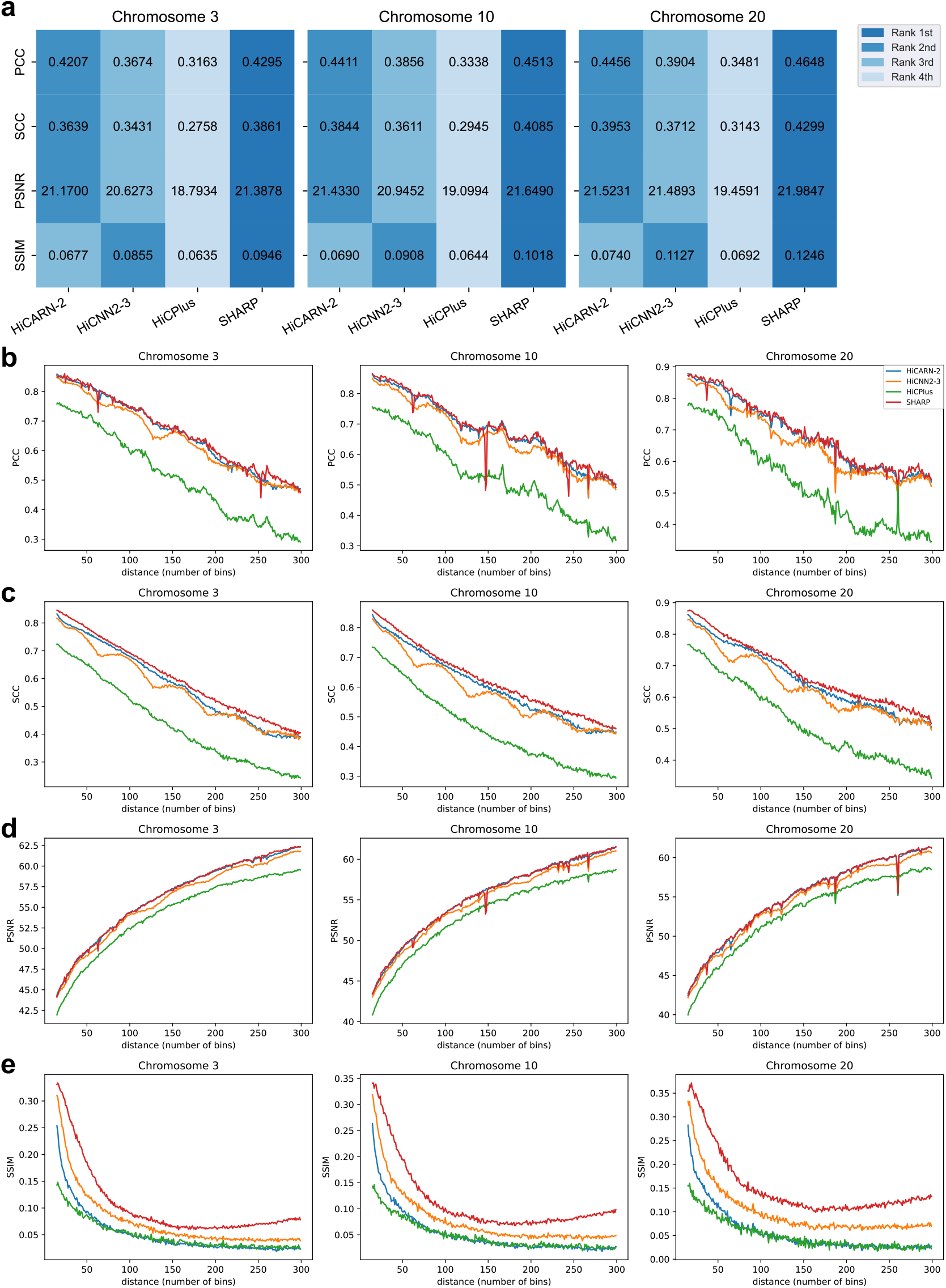
Resolution enhancement performance of SHARP as compared to three other methods. **a**, Performance of the different methods when applied to the left-out chromosomes based on four different performance measures. For all four measures, a larger value corresponds to better performance. **b-e**, Performance of the methods for region pairs at different genomic distances based on PCC (b), SCC (c), PSNR (d), and SSIM (e).

We further compared the performance of the four methods on pairs of genomic bins at different 1D distances separately. The results show that SHARP stably outperformed the other methods at almost all distance values for all the considered metrics (Figure 2b-e). HiCARN-2’s performance was close to SHARP in terms of PCC and PSNR, but it had low performance in terms of SSIM.

To assess the robustness of these results, we repeated the comparisons using 1) low-resolution contact matrices down-sampled from the high-resolution GM12878 matrix at other sampling rates (1/50 [*∼*50kb resolution] and 1/100 [*∼*90kb resolution], instead of the original 1/150 [*∼*200kb resolution]), 2) an actual low-resolution contact matrix from a GM12878 Hi-C data set with low sequencing depth (*∼*200kb resolution), instead of one down-sampled from a high-resolution matrix, and 3) high-resolution and down-sampled Hi-C data of another cell line (H1-hESC, instead of the original GM12878). The results show that SHARP also outperformed the other methods in most cases (Figure S1-S3).

### SHARP avoids creation of artificial structures

The four standard performance measures used in the comparisons above do not specifically quantify artifacts caused by the patches. We examined the resolution-enhanced contact matrices produced by the three existing methods and found artifacts at patch boundaries in all of them that do not appear in the actual high-resolution contact matrix (Figure 3a-e, Figures S4-S11). To systematically evaluate the prevalence of this issue, we invented the Normalized Boundary Difference (NBD) measure, which computes the average difference between values across a patch boundary normalized by the average difference between immediate neighboring values within the patch (Figure S12, Methods). A large NBD value indicates abrupt changes across patch boundaries that could be due to artifacts. Based on the NBD measure, SHARP produced a much lower level of artifacts than the other methods (Figure 3f).

**Figure 3:**
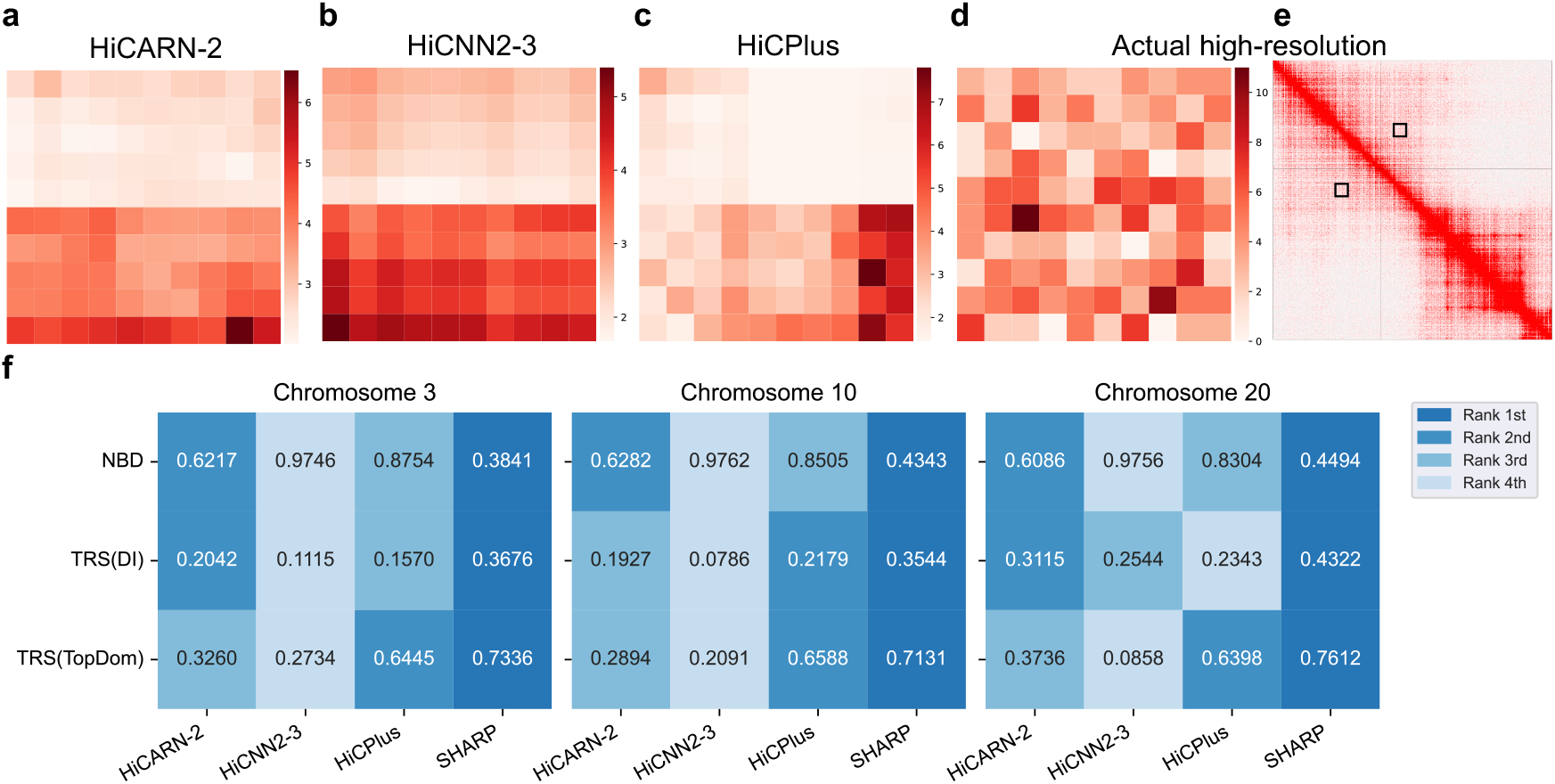
Artificial structures in resolution-enhanced Hi-C contact matrices and their avoidance by SHARP. **a-c**, An example area (Chr20:12.135Mb-12.185Mb *×* Chr20:12.69Mb-12.74Mb) of the resolution-enhanced contact matrix produced by HiCARN-2 (a), HiCNN2-3 (b), and HiCPlus (c). In all cases, the above and lower halves belong to two different patches. **d**, The actual high-resolution contact matrix of the same area. **e**, Location of the area in the whole high-resolution matrix of the chromosome marked by the black boxes. **f**, Performance of the different methods when applied to the left-out chromosomes based on three different measures. For the NBD measure, a smaller value corresponds to better avoidance of artificial structures; for the TRS measures, a larger value corresponds to better avoidance of artificial TADs.

To evaluate the downstream impacts of these artificial structures, we called TADs from the resolution-enhanced contact matrices and compared them with the TADs called from the actual high-resolution contact matrix. We defined the TAD-related score (TRS) to quantify how well artificial TADs caused by the patch boundaries are avoided (Methods). A large TRS value indicates better avoidance of these artificial TADs. Comparing the TRS values based on TADs called by two different methods (Directionality Index [DI]^22^ and TopDom^32^), SHARP produced fewer artificial TADs at the patch boundaries than the other three methods (Figure 3f).

Again, we repeated the comparisons with different down-sampling rates, an actual low-resolution contact matrix, and data from another cell line, which confirmed the robustness of SHARP in avoiding the creation of artificial structures at patch boundaries (Figure S13-S15).

To evaluate whether the signal decomposition stage of SHARP is important for avoiding artificial structures, we performed an ablation study that compared SHARP with a simplified version of it that does not perform signal decomposition but instead trains the MAXIM model using the contact matrix directly (“SHARP-NoDecomposition”, Methods). The results confirm that signal decomposition is useful for avoiding artificial structures (Figure S16). Interestingly, this simplified version of SHARP also outperformed the three existing methods in resolution enhancement performance (Figure S17), suggesting that the use of both local and global attention led to some performance improvements independent of signal decomposition.

### Identification of statistically significant interactions

An important use of resolution-enhanced Hi-C contact matrices is identifying high-resolution functional chromatin interactions. To evaluate SHARP’s ability to facilitate this, we started by comparing the statistically most significant interactions in the resolution-enhanced contact matrices with those in the actual high-resolution contact matrix (Methods). These interactions are the ones with the strongest enrichment of contact frequencies as compared to random expectation, and they represent promising candidates of functionally important interactions^33^. Based on two different evaluation measures, the significant interactions identified by FitHiC2^34^ produced by SHARP are most consistent with those identified from the actual high-resolution matrix no matter the whole reconstructed matrix was considered (Figure 4a-b) or only the imputed regions (Figure S18). SHARP also achieved the highest performance when applying the same analysis to the interactions identified by HiCCUPS^7^. However, the total number of interactions fluctuated considerably, making the results insufficiently robust.

**Figure 4:**
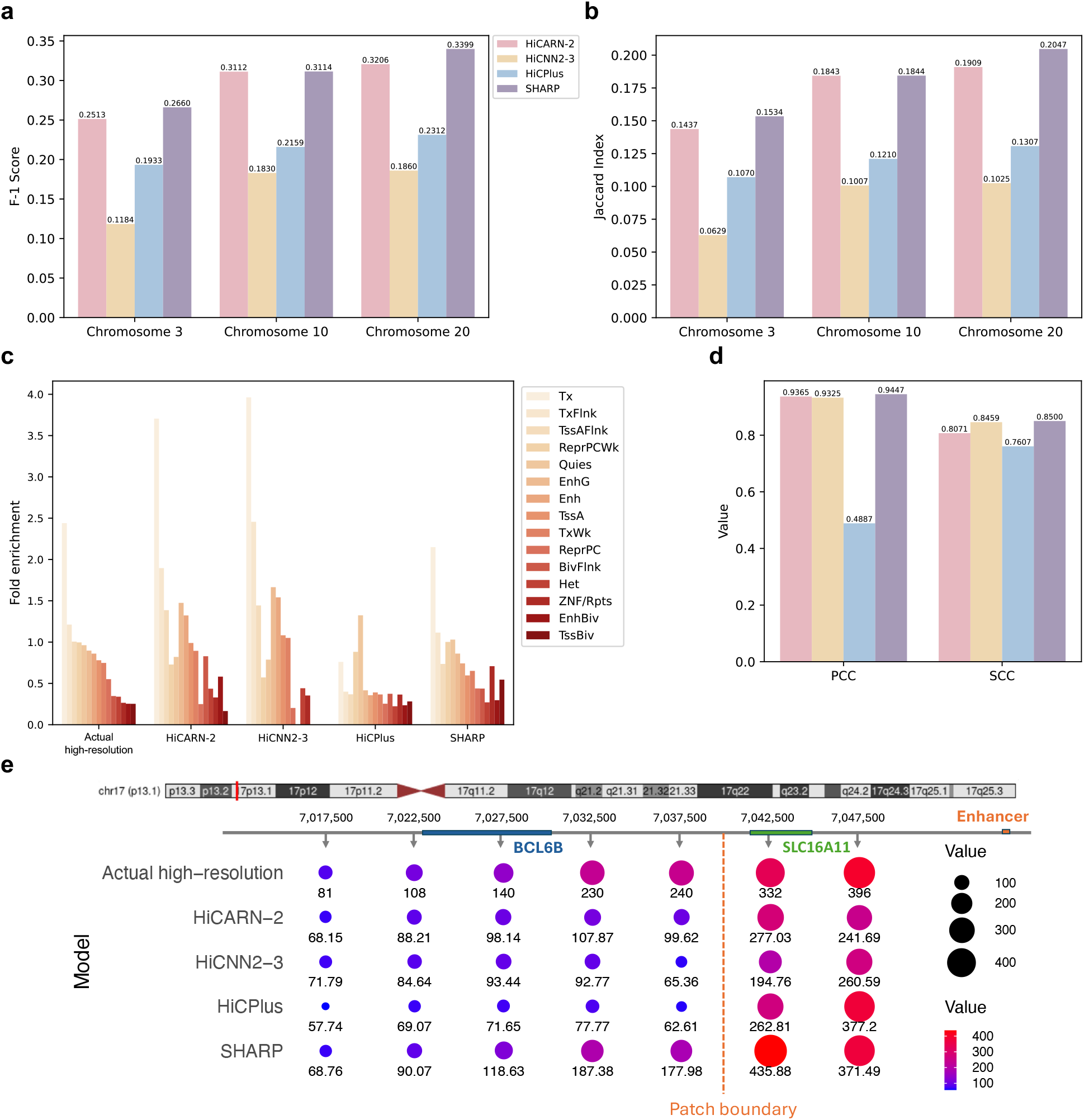
Significant chromatin interactions identified by SHARP as compared to three other methods. **a-b**, Comparing the statistically significant chromatin interactions identified from the resolution-enhanced contact matrices produced by the methods with those identified from the actual high-resolution matrix, quantified by the F-1 score (a) or Jaccard Index (b). **c**, Enrichment of the identified chromatin interactions at different chromatin states of Chr 20, including strong transcription (Tx), quiescent/low (Quies), weak repressed Polycomb (ReprPCWk), transcription at gene 5’ and 3’ (TxFlnk), weak transcription (TxWk), flanking active TSS (TssAFlnk), enhancers (Enh), genic enhancers (EnhG), active TSS (TssA), bivalent/poised TSS (TSSBiv), repressed Polycomb (ReprPC), flanking bivalent TSS/Enh (BivFlnk), ZNF genes + repeats (ZNF/Rpts), bivalent enhancer (En-hBiv), and heterochromatin (Het). The chromatin states are sorted in descending order of their enrichment values according to the actual high-resolution matrix. **d**, Correlation of the enrichment values with those computed based on significant interactions identified from the actual high-resolution matrix. **e**, Specific CRE-gene interactions identified by the different methods.

Next, we examined the chromatin states of loci involved in these significant interactions (Methods). Judged by fold enrichment values, all methods produced significant interactions in chromatin states resembling those obtained from the actual high-resolution contact matrix (Figure 4c, Figure S19a-b). However, some important differences are also observed. For example, in Chromosome 20, comparing the enrichment of the significant interactions in the strong transcription state (“Tx”), SHARP’s results are closest to the actual high-resolution matrix while the other methods produced enrichment values either too large (HiCARN-2 and HiCNN2-3) or too small (HiCPlus) (Figure 4c). Quantitatively, the enrichment values computed from the actual high-resolution Hi-C matrix are most consistently correlated with the enrichment values of SHARP (Figure 4d, Figure S19c-d).

We also used independently supported interactions between genes and cis-regulatory elements (CREs) to compare the significant chromatin interactions identified from the different resolution-enhanced contact matrices. Figure 4e shows an example of an enhancer (Chr17:7,130,223-7,134,160) cataloged in multiple databases (NCBI Reference Sequence ^35^, FANTOM5^36^, and ENCODE^37^) that interacts with two protein coding genes, *BCL6B* and *SLC16A11*^38^. Both interactions are supported by high gene association scores. In both the actual high-resolution Hi-C matrix and the resolution-enhanced matrices reconstructed by all four methods, the enhancer has the strongest interaction with *SLC16A11*. This is not surprising because *SLC16A11* is relatively close to the enhancer and they are both in the same patch defined by the resolution enhancement methods. However, only SHARP was able to infer a strong interaction between the enhancer and *BCL6B*, which reside in the same TAD and their interaction is supported by the actual high-resolution Hi-C matrix. The other three methods likely missed this interaction because the enhancer and the *BCL6B* locus were located in two different patches, which created difficulties for them to determine the interaction frequency accurately relative to the interaction frequency between the enhancer and the *SLC16A11* locus.

### Applicability of the human models to mouse data

In the actual use of a Hi-C resolution-enhancement method, it is given only a low-resolution matrix and has to directly enhance it without access to any high-resolution Hi-C data from the same sample. This requires applying a model previously trained on data from another sample. To assess such generalizability of the SHARP models, we performed model training using matched high- and low-resolution Hi-C contact matrices from human GM12878 cells and applied the models to enhance a low-resolution Hi-C matrix from mouse embryonic stem cells (mESCs). For benchmarking purposes, the same procedure was applied to other models, including HiCARN-2, HiCNN2-3, and HiCPlus. Chr 3, 10, and 19 were used as the test set in this analysis.

By having actual high-resolution data of the mESC contact matrix left-out from the training process, we were able to evaluate the performance objectively. In terms of resolution enhancement performance, SHARP performed the best based on SSIM and shared the top rank with HiCARN-2 based on the other three measures. Figure 5a-d). In terms of artificial structure avoidance performance, SHARP consistently outperformed the other three methods in all cases (Figure 5e).

**Figure 5:**
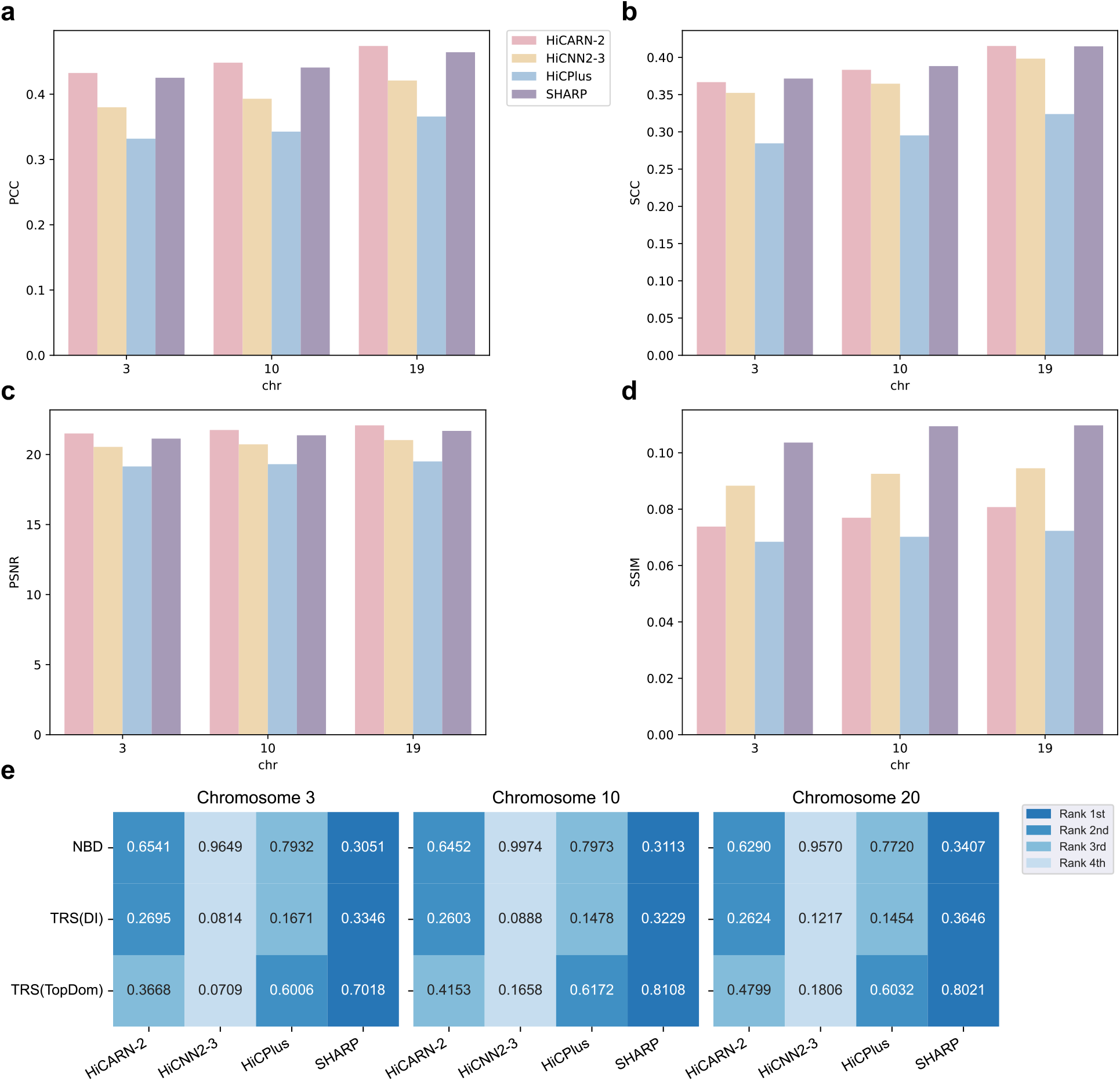
Performance of SHARP as compared to three other methods when applied to a Hi-C contact matrix of the mESC cell line. **a-d**, Resolution enhancement performance. **e**, Artificial structure avoidance performance.

### Fusing the resolution-enhanced contact matrices produced by SHARP and other methods

In computer vision, it has been shown that fusing the outputs of different models is an effective way to improve overall modeling performance since different models can complement each other^39–42^. To explore this possibility, we developed SHARP-Fusion, which uses supervised back-projection^43^ to integrate the enhanced Hi-C contact matrices produced by SHARP and other existing methods (Figure S20a, Methods). The back-projection procedure projects a base contact matrix onto another matrix, identifies the difference between the latter and the projection, and then adds the back-projected difference to the base matrix. This is performed multiple times to integrate information contained in the enhanced matrices produced by different existing methods into the enhanced matrix produced by SHARP. The projection is performed using convolutional layers and transposed convolutional layers, the parameters of which are learned using the chromosomes in the training set (Methods).

Comparing SHARP and SHARP-Fusion, we found that SHARP-Fusion generally performs better in terms of resolution enhancement accuracy (Figure S20b). However, it creates more artificial structures at the patch boundaries (Figure S20c), likely inherited from the contact matrices produced by the other three methods.

## Discussion

In this study, we have introduced SHARP that accurately enhances the resolution of Hi-C contact matrices while avoiding the creation of artificial structures at patch boundaries commonly created by other methods. Our main methodological innovations include 1) decomposition of the Hi-C contact matrices into three types of signals (due to 1D distance, contiguous domains, and other intricate structures, respectively) and enhance them separately, and 2) use of the both local and global attention mechanisms for enhancing the third type of signals. Using actual high-resolution Hi-C data of left-out chromosomes not involved in model training, we have shown that SHARP outperformed three other state-of-the-art methods in terms of both resolution enhancement performance and avoidance of creating artificial structures. We have also shown that these results are robust against 1) different ways to produce the low-resolution matrix (actual experimental data versus down-sampling of high-resolution data), 2) different down-sampling rates, and 3) data from different samples (training and testing both on GM12878 or both on h1-hESC) or another species (training on GM12878 and testing on mESC).

We have also performed an ablation study to show that the signal decomposition stage of SHARP is useful for avoiding the creation of artificial structures. At the same time, we have shown that the combined use of local and global attention is sufficient to improve resolution enhancement performance over existing methods even without signal decomposition.

In terms of computational resources required, SHARP required more CPU time but was on par with the other methods in terms of GPU time and memory usage (Table S1). Among the methods compared, HiCPlus had the lowest computational requirements but its resolution enhancement performance was usually lowest in our computational experiments.

Comparing SHARP and SHARP-Fusion, SHARP tends to create less artifacts at the patch boundaries while SHARP-Fusion tends to achieve better overall resolution enhancement performance. This suggests that the fusion procedure integrates some useful signals from the contact matrices produced by the other three methods that are missed by SHARP, but at the same time artificial structures in them are inadvertently also carried over. To combine the advantages of SHARP and SHARP-Fusion, one potential way is to use SHARP-Fusion for resolution enhancement but for all significant chromatin interactions and chromatin domains called, use the results of SHARP to confirm or reject them. Alternatively, the loss function and the overall model training procedure could be modified to take into account both enhancement performance and smoothness at patch boundaries.

SHARP and other Hi-C resolution enhancement methods are most useful when high-resolution data cannot be produced due to limited high-quality materials in the biological samples, such as patient biopsies. On the other hand, some of these samples also contain complex genomic rearrangements such as intra-chromosomal and inter-chromosomal translocation, which can create special patterns in the Hi-C data. Most existing Hi-C resolution enhancement methods have been mainly tested on data produced from cell lines that have abundant biological materials for the experiments, relatively low genomic complexity, and low heterogeneity among genomes in different cells. Further studies are needed to test whether these methods can also perform well on more difficult data. It may be useful to incorporate information from specialized tools such as structural variant callers designed for Hi-C data.

## Methods

### Details of the SHARP method

We model a Hi-C contact matrix by the summation of three types of signals, namely signals due to 1D proximity, signals due to within-domain and between-domain contacts of contiguous domains (such as TADs), and signals due to other fine structures. Here we describe how we isolate these three types of signals from a low-resolution matrix, enhance them separately, and sum them back to form a high-resolution matrix.

#### Modeling signals due to 1D proximity

As proposed previously^27^, the contact frequency between two genomic loci in a chromatin fiber separated by a genomic distance of *d* can be described by the following formula:

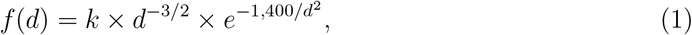

where *k* is a proportionality constant that reflects the efficiency of the cross-linking reaction.

When applying this formula to model contact frequencies in a Hi-C matrix, we discretized all genomic distances to multiples of the bin size.

We tested the goodness-of-fit of different functional classes for determining a suitable value of *k*. We found that the formula *k* = 7.0031 *× S*^0.7578^ provides a good starting point, where *S* is the sum of the whole matrix. We then fine-tuned *k* by testing different values around this number and finding the value that provided the best balance between removing signals on the diagonal and preserving signals in rectangles, aided by the Hi-C data visualization tool Juicebox^44^. Using this procedure, we found *k* = 3.5 *×* 10^6^ worked best for the low-resolution GM12878 matrix at subsampling rate of 1/150 (Figure S21) and *k* = 4 *×* 10^7^ worked best for the high-resolution GM12878 matrix (Figure S22).

#### Modeling signals due to contiguous domains

After subtracting out signals due to 1D proximity from the contact matrix, SHARP next identifies and subtracts out signals due to contiguous domains. These signals appear as rectangles that contain larger contact frequencies than surrounding areas (Figure S23). These rectangles can take the form of either squares that appear along the matrix diagonal, which contain interactions between different loci within the same domain, or off-diagonal stripes^45^, which contain interactions between loci from two different domains.

To cope with potentially very complex genome structures, we allow these rectangles to be nested and overlapping. Effects of overlapping rectangles sum together to produce the overall signals due to contiguous domains.

The detection of these rectangles somewhat resembles the task of TAD calling. While some current methods can identify hierarchical TAD structures or work on low-resolution data^46,47^, they are insufficient for our purpose due to our need for detecting off-diagonal structures and modeling effects of rectangles on contact frequencies. Therefore, we developed our own procedure for detecting the rectangular blocks and modeling their effects. Here we first give a high-level description of the procedure, and then provide further details.

#### Overview of the procedure

Our procedure detects and subtracts out the rectangles in the following order:

1. Detect all on-diagonal rectangles.
2. Subtract out all on-diagonal rectangles.
3. Detect and subtract out an off-diagonal rectangle.
4. Repeat #3 until no more off-diagonal rectangles can be detected.

On-diagonal and off-diagonal rectangles are handled differently because on-diagonal rectangles overlap with each other much more frequently than off-diagonal rectangles. Therefore, if an on-diagonal rectangle is subtracted out immediately after its detection before knowing the presence of other overlapping rectangles, its effects on contact frequencies may not be modeled accurately. In contrast, since off-diagonal rectangles seldom overlap with each other, they can be detected and subtracted out independently using a simpler procedure.

To prevent our procedure from being misled by problematic regions that possibly contain spurious signals, we excluded all bins (and moved the following bins up) that overlap the low mappability regions and high signal regions on the ENCODE blacklist^48^, obtained from https://github.com/Boyle-Lab/Blacklist/blob/master/lists/hg38-blacklist.v2.bed.gz.

#### Detecting on-diagonal rectangles

The procedure for detecting on-diagonal rectangles is shown in Figure S24. Each on-diagonal rectangle is a square fully specified by its starting and ending bins. SHARP searches for on-diagonal rectangles by considering each bin as a potential starting bin in turn. From a potential starting bin, SHARP defines a thin stripe as the potential upper edge of the rectangle. The average contact frequency within the stripe is compared to that of an area of the same length immediately outside the tentative rectangle, which serves as reference. If the former has a larger average frequency, the tentative rectangle will be recorded and further refined in the next stage. To detect rectangles of different sizes, several stripe lengths are tested for each potential starting bin.

For each of these tentative rectangles, SHARP then attempts to expand it by comparing the average contact frequencies within the rectangle, the expansion, and the area immediately outside. The expansion process is then repeated iteratively until the next expansion would not increase the average contact frequency of the whole rectangle.

Details of the whole procedure is given in Algorithm 1.

#### Categorizing on-diagonal rectangles

While a large proportion of contact frequencies within an on-diagonal rectangle are larger than those in the immediate surrounding areas, there are three different patterns commonly observed, namely fairly uniform signals in the whole rectangle (Figure S25a), stronger signals in the lower triangle (Figure S25b), and stronger signals in the upper triangle (Figure S25c). The first category includes domains in which most loci can be freely in contact with each other, while the second and third categories include domains that have loci close to one domain boundary more flexibly interacting with other loci in the domain than those close to the other domain boundary.

Based on this observation, for each on-diagonal rectangle detected, SHARP puts it into:

##### Algorithm 1 Detecting on-diagonal rectangles

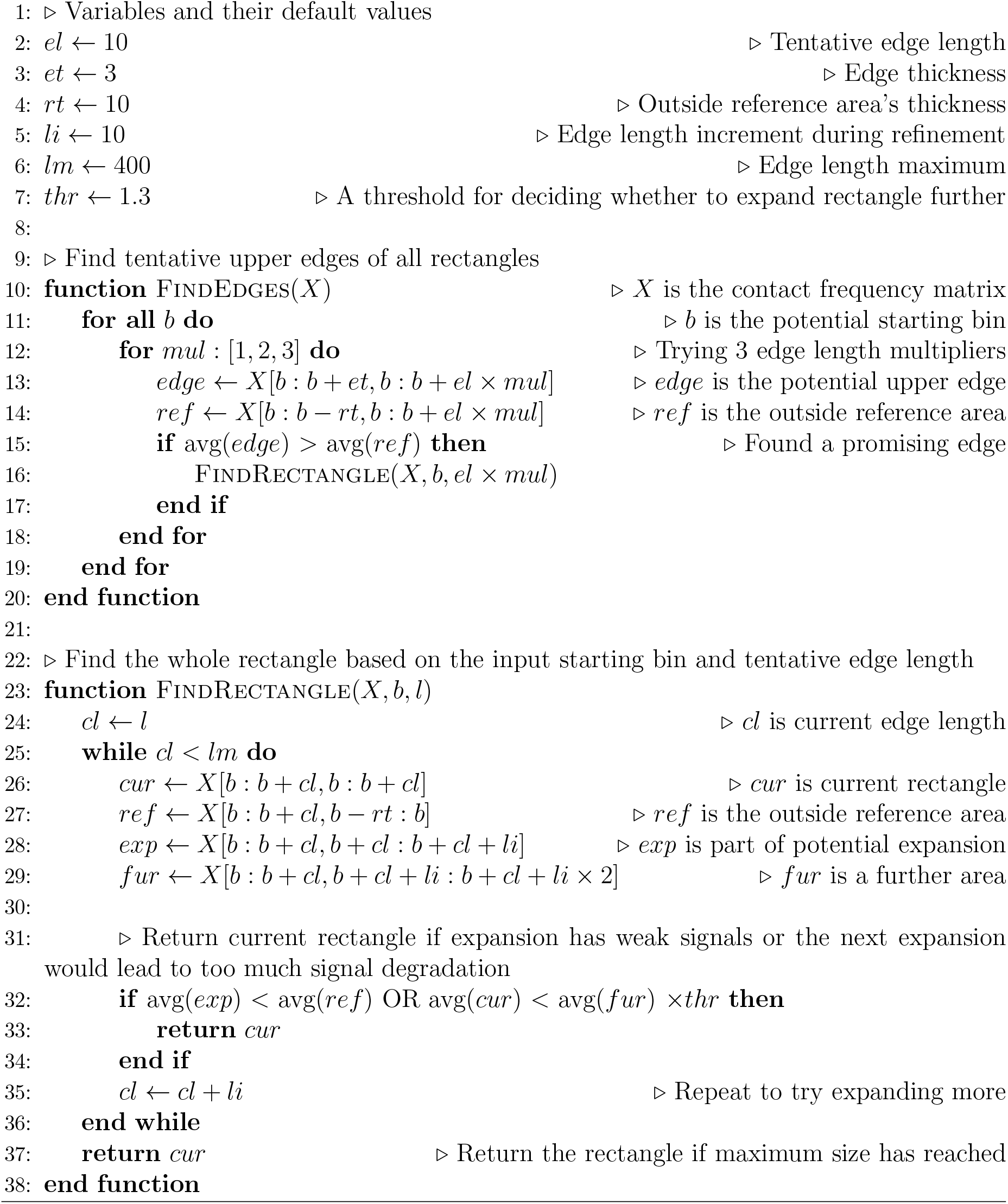

- The upper triangle category if the average contact frequency in the upper triangle is *>*1.5 times the average in the lower triangle
- The lower triangle category if the average contact frequency in the lower triangle is *>*1.5 times the average in the upper triangle
- The uniform signal category otherwise

#### Modeling and subtracting out effects of on-diagonal rectangles

For all three categories of on-diagonal rectangles, SHARP models its effects by a constant multiplying factor *m* that is obtained by comparing strong values within the rectangle with some baseline values. Where the strong values and baseline values are obtained differs for the three categories (Figure S26):

- For the uniform signal category, we take contact frequencies in the whole rectangle as the strong values and contact frequencies in the immediate outside areas as the baseline values.
- For the lower triangle category, we take contact frequencies in the lower triangle as the strong values and contact frequencies in the upper triangle as the baseline values.
- For the upper triangle category, we take contact frequencies in the upper triangle as the strong values and contact frequencies in the lower triangle as the baseline values.

Let *μ* and *μ*^*′*^ be the average of the strong values and the average of the baseline values, respectively. The multiplying factor *m* is computed as:

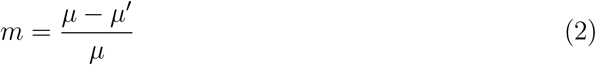

To subtract out the signals due to the rectangle, each contact frequency *x* within the “strong value” area is adjusted using the multiplying factor:

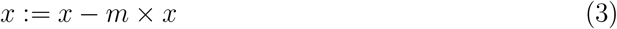

In other words, each original contact frequency in the rectangle is 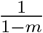 times of the adjusted value.

After this adjustment, for each on-diagonal rectangle in the uniform signal category, the effect of the whole rectangle is expected to be subtracted out such that contact frequencies inside and immediately outside the rectangle should be similar. On the other hand, for the other two categories, the procedure only turns the rectangle into one that has uniform signals but it may still have stronger signals than the immediate outside areas. Therefore, for each rectangle originally in these categories, we apply this adjustment one more time but treating it as in the uniform signal category during this second round of adjustment.

Since different rectangles can be nested or overlapping, SHARP performs the above operations on small rectangles first before large rectangles, by ordering all rectangles by their sizes. By doing so, if a small rectangle is within a big nested rectangle, the baseline values taken immediately outside the small rectangle will likely be still within the big rectangle. As a result, when the small rectangle is processed, only the effect due to it will be adjusted while the effect due to the big rectangle will remain, which will be subtracted out when it is the big rectangle’s turn to be processed.

#### Detecting off-diagonal rectangles and modeling and subtracting out their effects

Off-diagonal rectangles are detected in a way similar to on-diagonal rectangles, except that 1) they can be anywhere in the matrix rather than only along the diagonal and 2) they are not necessarily squares. Therefore, each off-diagonal rectangle needs to be specified by two pairs of starting and ending bins instead of just one pair. During the searching process, when the upper edge of a potential rectangle is identified, both the height and width of the rectangle need to be determined by applying the same procedure to them separately.

When the whole rectangle is defined, its effect on contact frequencies is modeled and subtracted out using the same way as an on-diagonal rectangle that belongs to the uniform signal category.

#### Modeling other signals

After signals due to 1D proximity and contiguous domains have been subtracted out from the contact matrix, SHARP next models relationships between the remaining signals in the low-resolution and high-resolution matrices using deep learning. These remaining signals are mostly due to fine-grained local structures, and therefore we can use the standard strategy of dividing up the contact matrix into regular patches. We chose MAXIM^28^ as the model architecture, which uses the attention mechanism of the highly successful transformers^49,50^. Overcoming limitations of full self-attention^51,52^, which can only be applied to small patches, and local attention^53,54^, which leads to a small receptive field, MAXIM combines both local and global attention.

MAXIM achieves this using a special encoder-decoder architecture (Figure S27a). A conventional autoencoder aims at reconstructing the input from the embedding captured by the bottleneck layer. MAXIM modifies this objective by considering both the original input and down-sampled versions of it, which ignore some details but instead focus on higher-level information. Each version of the data has its own encoder-decoder pair, but outputs of the different encoders are integrated using cross-gating blocks (GCBs), thereby allowing features from the different versions of the data to jointly reconstruct each version of it.

Each encoder, decoder, and bottleneck block is composed of layers of multi-axis gated multi-layer perceptron (MLP) and residual channel attention blocks^55,56^ (Figure S27b). The multi-axis gated MLP block performs both regional and dilated attention. Regional attention captures local information more effectively and follows the principles of convolutional operations, while dilated attention is primarily used to fuse long-range features (i.e., global feature fusion) and is akin to dilated convolutional operations. The residual channel attention block adaptively recalibrates channel-wise features to emphasize on the most crucial information.

The CGBs were designed to enhance feature propagation in skip-connections by selectively gating features through contextual information (Figure S27c). At the core of the CGB is the multi-axis cross-gating mechanism, which conditions one feature map on another by applying regional and dilated attention, akin to the Multi-axis Gated MLP block. Three CGBs are employed in the overall MAXIM architecture (Figure S27a), each corresponding to a different input scale to capture contextual interactions at various stages of feature extraction and reconstruction.

In our adoption of MAXIM, we use Mean Absolute Error (MAE)^57^ as the loss function since it is relatively robust to outliers and does not penalize large error terms as strongly as some other loss functions such as the Mean Squared Error (MSE). In Hi-C contact matrices, there are occasionally some extremely large values and it is important to prevent the training process from focusing on modeling these values.

To train the MAXIM model, we used the Adam optimizer^58^ with an initial learning rate of 2 *×* 10^*−*4^. This learning rate was gradually decreased to 10^*−*7^ using cosine annealing decay.

#### The complete training process

The overall model training process of SHARP requires paired low-resolution and highresolution Hi-C contact matrices of the same sample (Figure 1g).

First, signals due to 1D proximity are subtracted out from both the low-resolution and high-resolution matrices independently.

Next, from the resulting low-resolution matrix, on-diagonal and off-diagonal rectangles are detected, modeled, and subtracted out. For each rectangle, three types of parameters specific to it are recorded, namely 1) its starting and ending bins (one pair for an on-diagonal rectangle, two pairs for an off-diagonal rectangle) 2) its rectangle category, and 3) multiplying factor *m*. These results are then applied to the high-resolution matrix to subtract out signals due to contiguous domains, using the parameters learned from the low-resolution matrix directly.

Finally, taking the remaining signals from the low-resolution and high-resolution matrices together, the MAXIM model is trained to minimize the error in enhancing the low-resolution one to become the high-resolution one.

#### Applying SHARP to enhance a low-resolution Hi-C contact matrix

In the actual use of SHARP, it is given only a low-resolution Hi-C contact matrix as input and is tasked to reconstruct a high-resolution of it using the model previously learned from another sample. The procedure involves several steps:

1. Signals due to 1D proximity are subtracted out from the low-resolution matrix.
2. Signals due to contiguous domains are detected, modeled, and subtracted out from the low-resolution matrix.
3. The previously learned MAXIM model is applied to the resulting low-resolution matrix to reconstruct the high-resolution matrix without the first two types of signals.
4. Signals due to contiguous domains are added to the reconstructed high-resolution matrix using the same parameters obtained in Step 2.
5. Signals due to 1D proximity are added to the reconstructed high-resolution matrix using a proportionality constant obtained from the high-resolution matrix involved in the original model training of SHARP.

In the reconstructed high-resolution matrix, if there are any negative values, they are all set to zero. Also, for values larger than the maximum value of the low-resolution contact matrix multiplied by a threshold, we set them to the median of their surrounding four values. The default value of this threshold is the ratio of total contact frequency of the high-resolution matrix to that of the low-resolution matrix of the training data.

### Other Hi-C resolution enhancement methods compared

To assess the performance SHARP, we compared it with three state-of-the-art Hi-C resolution enhancement methods, HiCPlus^8^, HiCARN^19^, and HiCNN2^25^.

In the original publications of HiCARN and HiCNN2, multiple variants of these models are described. We chose the best-performing ones based on the results presented in the original publications, namely HiCARN-2 for HiCARN and HiCNN2-3 for HiCNN2.

When using these methods, we always set their parameters following recommendations in the original publications.

### Details of the SHARP-Fusion method

The back-projection method^43^ used by SHARP-Fusion was originally developed for image enhancement, with a goal of reconstructing a high-resolution image by combining multiple low-resolution images sequentially. In our adoption of the method, we used it to reconstruct a high-resolution Hi-C contact matrix by combining multiple reconstructed high-resolution matrices produced by different methods sequentially (Figure S20a).

Specifically, the output (i.e., reconstructed high-resolution matrix) of SHARP is used as the base matrix. Information contained in the output of HiCPlus that is useful for reconstructing the high-resolution matrix but missing in the base matrix is added to it. The same procedure is then repeated to add useful information from the outputs of HiCARN-2 and HiCNN2-3 sequentially. Mathematically, this process can be summarized by the following formulas:

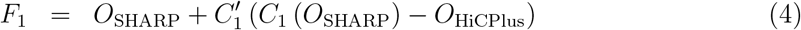

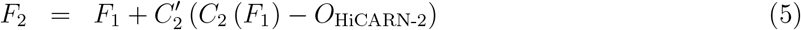

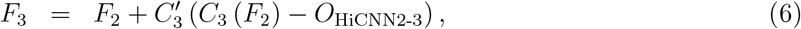

where *O*_X_ is the output of method X, *C*_*i*_ and 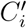 are convolutional layers for projecting one matrix onto another and projecting the difference back, respectively, in the *i*-th sequential fusion stages, and *F*_*i*_ is the fusion result of the *i*-th stage. The final output, *F*_3_, is the output of SHARP-Fusion.

During model training, parameters in the 6 convolutional layers are learned together to minimize reconstruction errors as compared to the actual high-resolution contact matrix. To train the model, we used the Adam optimizer with a learning rate of 5 *×* 10^*−*4^.

When using SHARP-Fusion to enhance a low-resolution matrix without paired highresolution data, the above formulas are applied directly using the learned parameters.

### Data sets used to compare the different methods

#### Data sources

All the Hi-C data used in this study were produced by *in situ* Hi-C^7^. We downloaded the data from the 4D Nucleome Data Portal^2,59^, with experiment set accession numbers 4DNES3JX38V5 (high-resolution GM12878), 4DNESBJ1KYYH (low-resolution GM12878), 4DNESX75DD7R (H1-hESC), and 4DNESDXUWBD9 (mESC).

#### Data processing

For each Hi-C data set, we converted it into a symmetric contact matrix with 5kb bin size using Juicer^60^ without applying any normalization. We did not apply normalization because some of the methods require unnormalized data, and some previous methods for enhancing Hi-C resolution were also tested on unnormalized data^8,19^. Each entry of the matrix thus contains the interaction frequency between a pair of genomic bins.

Since in our computational experiments our goal was to infer high-resolution Hi-C matrices at 5kb resolution, for all Hi-C matrices we used, including the low-resolution and high-resolution ones, we used 5kb as the bin size.

We found some outlier values in the Hi-C data. For example, in the high-resolution GM12878 data, the largest contact frequency, between two bins on chromosome 16, has a value of 106,901, while in other chromosomes, the largest contact frequency is only around 2,000. These outlier values were likely caused by technical errors^61^. To avoid these outlier values from biasing the models, we set them to a more reasonably large value. Specifically, we sorted all contact frequencies in ascending order and identified the first value in the sorted list that is 10 or more larger than its previous value. That previous value was defined as the new maximum and all values after it on the sorted list were set to this maximum.

When we applied the models learned from GM12878 to enhance the low-resolution matrix of mESC, an additional scaling step was required because their contact frequencies had very different value ranges. We first computed the median of contact frequencies on the main diagonal in each of the two matrices. We then took the GM12878-to-mESC ratio of these medians and multiply every value in the mESC matrix by this scaling factor.

All Hi-C resolution enhancement methods received the same processed data as input.

#### Down-sampling for simulating low-resolution matrices

From an actual high-resolution Hi-C contact matrix, we performed down-sampling to simulate a low-resolution contact matrix of the same sample. For a down-sampling rate of *r*, for each contact in the high-resolution matrix, we kept it in the simulated low-resolution matrix with a probability of *r*. We implemented it by down-sampling each matrix entry in turn. For a matrix entry originally having a contact count of *c* in the high-resolution matrix, we sampled a value from the binomial distribution with mean *cr* and variance *cr*(1 *− r*) to become the corresponding contact count in the simulated low-resolution matrix.

### Experimental settings

#### Bin pairs for model training, selection, and evaluation

To enable unbiased performance evaluation, following previous studies^17,25^, we divided all chromosomes into three disjoint subsets. For human data (GM12878 and H1-hESC), Chromosomes 1, 2, 4-9, 11-15, 18, and 19 were used for model training. Some methods produced different possible models during the training process, such as the different MAXIM models produced by SHARP in different training epochs. The performance of these different models was compared using Chromosomes 16, 17, 21, and 22, and the one with the best performance (quantified by PCC *×* SCC after confirming they were both positive) was chosen. Finally, the model was evaluated using Chromosomes 3, 10, and 20, which were left-out from all these previous steps. These three chromosomes serve as representatives of long, medium, and short chromosomes, respectively.

When applying models learned from GM12878 to enhance the mESC contact matrix, mouse Chromosomes 3, 10, and 19 were used for performance evaluations.

For all the resolution enhancement methods, we used a patch size of 64 bins *×* 64 bins, i.e., 320kb *×* 320kb. Since there are relatively few very long-range chromatin contacts, following previous studies^8,17,29^, we considered only bin pairs that are no more than a certain genomic distance apart (set to 1.92Mb in this study) on the same chromosome. These bin pairs will be described as those included in the “evaluation set” below. For the remaining regions in the Hi-C contact maps, we directly copied the data from the low-resolution matrix to the inferred high-resolution matrix.

#### Default setting

In the empirical comparisons, by default the low-resolution contact matrix used was the one down-sampled from the corresponding high-resolution contact matrix at a sampling rate of 1/150 because it matches the resolution of the actual low-resolution GM12878 matrix. Accordingly, by default the “ground truth” high-resolution matrix used for performance evaluation was that actual high-resolution contact matrix.

In all the results reported, except for the changes explicitly stated (e.g., using a different down-sampling rate or using another sample), all other configurations remain the same as this default setting.

#### Alternative settings for robustness evaluation

In addition to the default setting, we also compared the performance of SHARP with the three other methods using some alternative settings. These include using other down-sampling rates (1/50 and 1/100) to simulate a low-resolution GM12878 contact matrix, using an actual low-resolution GM12878 contact matrix, and using Hi-C data from H1-hESC (with the low-resolution contact matrix of it produced by down-sampling at the rate of 1/150). In each of these cases, the paired high- and low-resolution matrices were used to perform training, model selection, and testing, based on the different sets of chromosomes as in the default setting.

#### Ablation study

To evaluate the significance of signal decomposition, we produced a variation of SHARP (called “SHARP-NoDecomposition”) that does not subtract out the first two types of signals. Instead, it takes the low-resolution matrix and directly trains a MAXIM model to enhance it.

### Evaluation measures for resolution enhancement performance

We used four standard evaluation measures to quantify the resolution enhancement performance of the methods compared. These measures consider different aspects of the performance and were commonly used in previous studies^17,19,29^. For each measure, we first computed its value in each patch, and then reported the average of these values across all patches in the evaluation set.

The first measure is Pearson correlation coefficient (PCC)^62^, which measures the linear correlation between two lists of values of the same length. PCC ranges from -1 to 1, with a larger value corresponding to a more positive linear relationship between the two lists. To use PCC to evaluate resolution enhancement performance, we took all contact frequencies in a patch in the reconstructed and actual high-resolution contact matrices ordered in the same way according to genomic locations of the defining bins.

The second measure is Spearman correlation coefficient (SCC)^63^. It is similar to PCC but instead of considering the contact frequencies directly, SCC considers their positions in the sorted list of contact frequencies. We used the stats.spearmanr function in the SciPy library, which gives all values in tie their average rank in the group. As compared to PCC, SCC emphasizes more on the order of the values than the exact values. As a result, SCC is less affected by outliers than PCC. On the other hand, SCC is more affected than PCC when there is a lot of similar values the order of which is not important.

The third measure is peak signal-to-noise ratio (PSNR)^64^. It compares the maximum value and the root mean square error between the reconstructed and actual high-resolution matrices. A larger value of PSNR indicates better reconstruction performance.

The fourth measure is structural similarity index measure (SSIM)^65^. It uses spatial dependencies to model structural features and compares these features in the reconstructed and actual high-resolution matrices.

### Evaluation measures for artificial structure avoidance performance

We proposed two measures for quantifying the performance of different resolution enhancement methods in avoiding the creation of artificial structures due to patch boundaries.

The first measure is Normalized Boundary Difference (NBD). Consider the left boundary of a patch *i* in a reconstructed high-resolution matrix. Let *x*^(*i*)^, *y*^(*i*)^, and *z*^(*i*)^ be the first column inside the patch, second column inside the patch, and first column outside the patch, respectively (Figure S12). If no artificial structures are created at this boundary, the internal difference between *x*^(*i*)^ and *y*^(*i*)^ should be similar to the cross-boundary difference between *x*^(*i*)^ and *z*^(*i*)^. Based on this, we defined the following measure considering the left boundaries of all patches together:

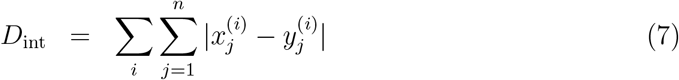

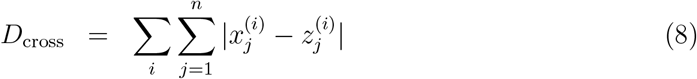

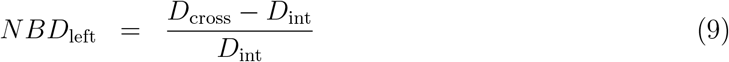

We also defined the same measure for the other three boundaries of the patches and used their average as the final performance measure:

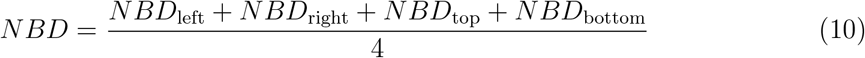

The second measure is TAD-related score (TRS). It quantifies incorrect TADs called from a reconstructed high-resolution matrix caused by the patch boundaries. Roughly speaking, a TAD called from a reconstructed matrix is considered incorrect if it is not also called from the actual matrix. However, there are three difficulties in defining such a quantitative measure. First, if a TAD is considered correct only if there is also a TAD called from the actual high-resolution matrix with exactly the same starting and ending bins as it, most TADs would be considered incorrect since TAD boundaries can be easily shifted by small changes of the contact matrix. Second, even if the definition of a correct TAD tolerates some shifts of the TAD boundaries, it is still not trivial to determine how TADs called from two matrices should be matched. Third, the number of TADs called can easily bias the results. For example, if from a reconstructed matrix very few TADs are called, the ratio of them with a starting bin or ending bin close to a patch boundary could be small just by chance, but this does not mean the resolution enhancement method used to reconstruct the matrix is unaffected by patch boundaries.

Taking all these into account, our TRS measure first considers the ratio of TADs called from the reconstructed matrix that have a starting bin or ending bin near a patch boundary. A larger ratio means potentially more incorrect TADs caused by the patch boundaries. However, some TADs called from the actual high-resolution matrix can also have starting or ending bins near patch boundaries. Therefore, our TRS measure further checks to what extent these actual TADs overlap the TADs called from the reconstructed matrix. A larger overlap means more support that the TAD boundaries close to patch boundaries are actually correct. Finally, these two components are combined to become the overall TRS measure.

Specifically, the procedure for computing TRS includes the following steps (more details of individual steps will be provided later):

1. Perform a matching of all the TADs called from the actual high-resolution contact matrix and all the TADs called from the reconstructed matrix. For each pair of overlapping TADs, an overlap ratio is computed based on their starting and ending bins. If a TAD cannot be matched, its overlapping ratio is 0.
2. Collect all TADs called from the actual high-resolution matrix that have a starting bin or ending bin near a patch boundary.
3. Compute the average overlap ratio of all the TADs collected in Step 2.
4. Compute TRS as *{*Average overlap ratio from Step 3*}* - *{*#TADs called from reconstructed matrix with starting or ending bin near a patch boundary*}* / *{*#TADs called from reconstructed matrix*}*

Based on this definition, a larger TRS value indicates a smaller number of incorrect TADs caused by the patch boundaries. If very few TADs are called from the reconstructed matrix, the average overlap ratio would be small since most TADs called from the actual matrix would be unmatched, and therefore the TRS value would not be large in that case.

The TAD matching is performed using a greedy algorithm. For each TAD *i* called from the actual high-resolution matrix and each TAD *j* called from the reconstructed matrix, an overlap ratio is computed as 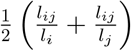, where *l*_*i*_ and *l*_*j*_ are the lengths (i.e., number of bins) of TAD *i* and TAD *j*, respectively, and *l*_*ij*_ is length of their intersection. All these overlap ratios are then sorted in descending order. The two TADs that give the largest overlap ratio are matched, and all overlap ratios of these TADs are removed from the sorted list. This matching process is then repeated until the largest overlap ratio is 0, at which time all remaining TADs are considered unmatched and they all have an overlap ratio of 0.

In Steps 2 and 5, the starting bin or ending bin of a TAD is considered near a patch boundary if they are differed by no more than 2 bins.

### Analysis of significant interactions

#### Definition of significant interactions

From a (reconstructed or actual) high-resolution Hi-C contact matrix, we applied FitHiC2^34^ and HiCCUPS^7^ to estimate the statistical significance of the contact frequencies. In FitHiC2, all bin pairs at a genomic distance between 15kb and 1Mb with a q-value less than 1 *×* 10^*−*7^ were considered to have a significant interaction. We utilized the default parameters specified by HiCCUPs for processing high-resolution Hi-C contact maps.

#### Evaluation measures for significant interactions

We used two common measures to evaluate the concordance between the significant interactions identified from the reconstructed and actual high-resolution matrices.

The first measure is the F-1 score. Let *R* and *A* be the sets of significant interactions identified from the reconstructed and actual high-resolution matrices, respectively, and |*X*| be the number of significant interactions in a set *X*. Precision computes the proportion of significant interactions identified from the reconstructed matrix that are also supported by the actual matrix; recall computes the proportion of significant interactions identified from the actual matrix that are also identified from the reconstructed matrix; finally the F-1 score aggregates these two measure by taking their harmony mean:

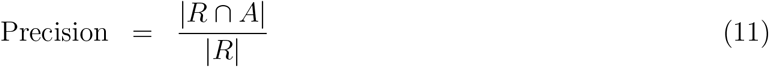

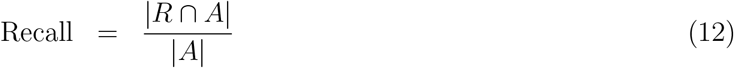

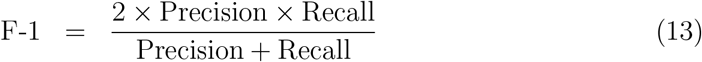

The second measure is the Jaccard index. It uses a different way to penalize both wrong and missing significant interactions by considering the number of interactions in the union of the two sets:

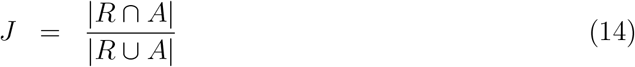

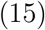

Both measures have a range between 0 and 1, with a larger value corresponding to better concordance between the two sets of significant interactions.

#### Analysis of chromatin states

We obtained the chromatin states of individual 200bp regions in the GM12878 genome inferred by a ChromHMM^66^ 15-state model lifted to hg38 from https://egg2.wustl.edu/roadmap/data/byFileType/chromhmmSegmentations/ChmmModels/coreMarks/jointModel/final/ produced by the NIH Roadmap Epigenomics Mapping Consortium^67^. For each genomic bin involved in a significant interaction, we recorded the number of base pairs in it that overlap with each chromatin state. By summing these base pair counts over the genomic bins, we computed the fraction of total genomic span of these bins that belongs to each chromatin state. Similarly, we computed the fraction of genomic span that belongs to this chromatin state in the whole genome. The ratio of the former to the latter was defined as the fold enrichment.

### Genomic distance-stratified analysis

In addition to computing overall performance values by considering all bin pairs included in the evaluation set together, we also computed performance values for bin pairs at different genomic distances apart separately.

For each of PCC, SCC, PSNR, and SSIM, for each genomic distance (i.e., number of genomic bins), we computed a single performance value using all contact frequencies in the reconstructed and actual high-resolution matrices of bin pairs separated exactly by that distance. In this case, the input data type for PSNR and SSIM was a one-dimensional vector of contact frequencies, rather than a two-dimensional sub-matrix from a patch as in the overall performance calculations.

## Code availability

The source code of SHARP and SHARP-Fusion and the trained models are available at https://github.com/Yip-Lab/SHARP.

## Acknowledgments

QC is supported by National Natural Science Foundation of China under Award Number 32100515 and CUHK direct grant for research under Award Numbers 2021.061 and 2022.080. KYY is supported by National Cancer Institute of the National Institutes of Health under Award Numbers P30CA030199 and R01CA287114, National Institute on Aging of the National Institutes of Health under Award Numbers R01AG085498 and U54AG079758, and internal grants of Sanford Burnham Prebys Medical Discovery Institute. The content is solely the responsibility of the authors and does not necessarily represent the official views of the National Institutes of Health.

## Contributions

KYY conceived the study. QL and KYY designed the methods. QL implemented the methods, collected and processed the data, and performed the data analysis. QL, KYL, CN, PLP, QC, and KYY interpreted the results. QL, QC, and KYY wrote the manuscript. All authors read and approved the final version of the manuscript.

## Supplementary information

### Supplementary Tables

**Supplementary Table S1:** Comparison of computational resources required by the different methods for one batch of training in the first epoch.

| Method | CPU Memory (MB) | GPU Memory (MB) | CPU Time (ms) | GPU Time (s) |
| --- | --- | --- | --- | --- |
| HiCARN-2 | 2322.85 | 666.85 | 788.17 | 2.62 |
| HiCNN2-3 | 2290.74 | 47.18 | 773.32 | 3.03 |
| HiCPlus | 2259.38 | 1.46 | 387.80 | 1.53 |
| SHARP | 2370.04 | 77.61 | 1290.00 | 3.01 |

### Supplementary Figures

**Supplementary Figure S1:**
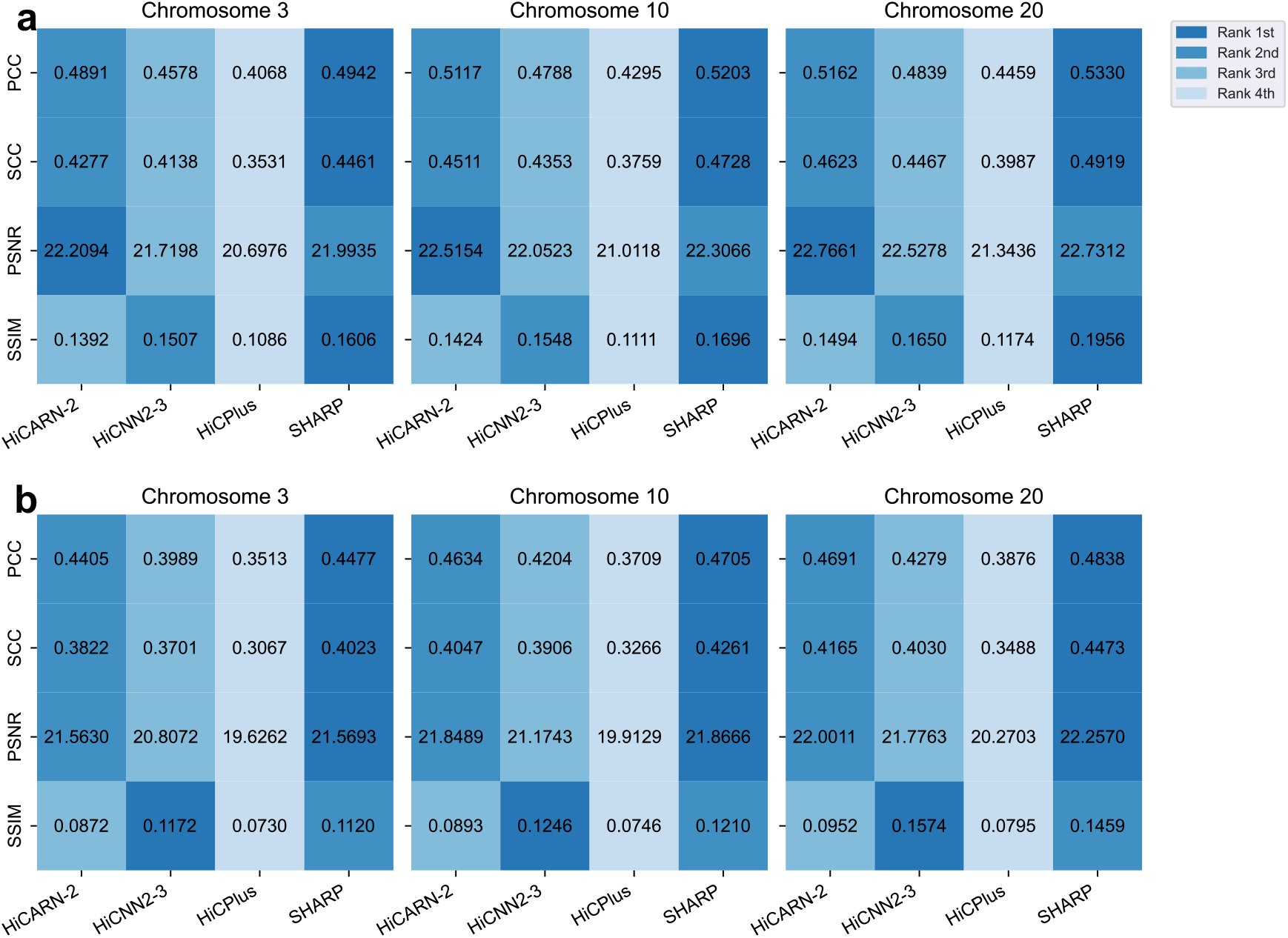
Resolution enhancement performance of SHARP as compared to three other methods when applied to low-resolution contact matrices down-sampled from a high-resolution matrix at different sampling rates. **a**, Sampling rate of 1/50. **b**, Sampling rate of 1/100.

**Supplementary Figure S2:**
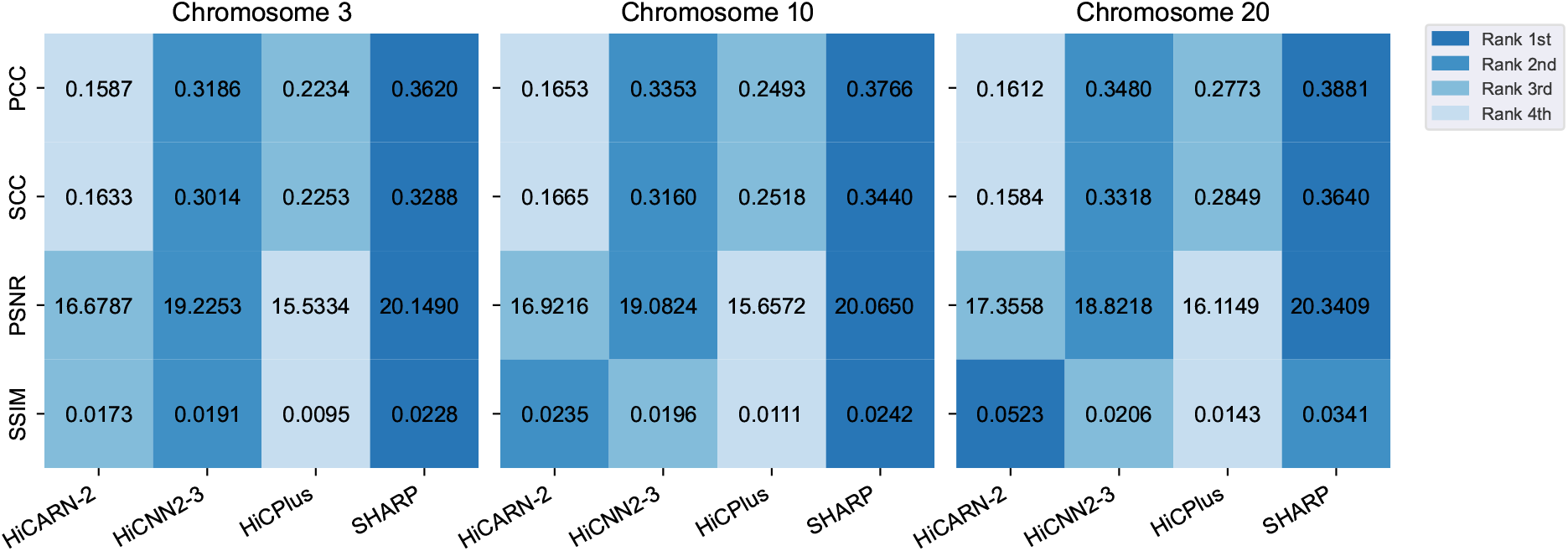
Resolution enhancement performance of SHARP as compared to three other methods when applied to an actual low-resolution contact matrix.

**Supplementary Figure S3:**
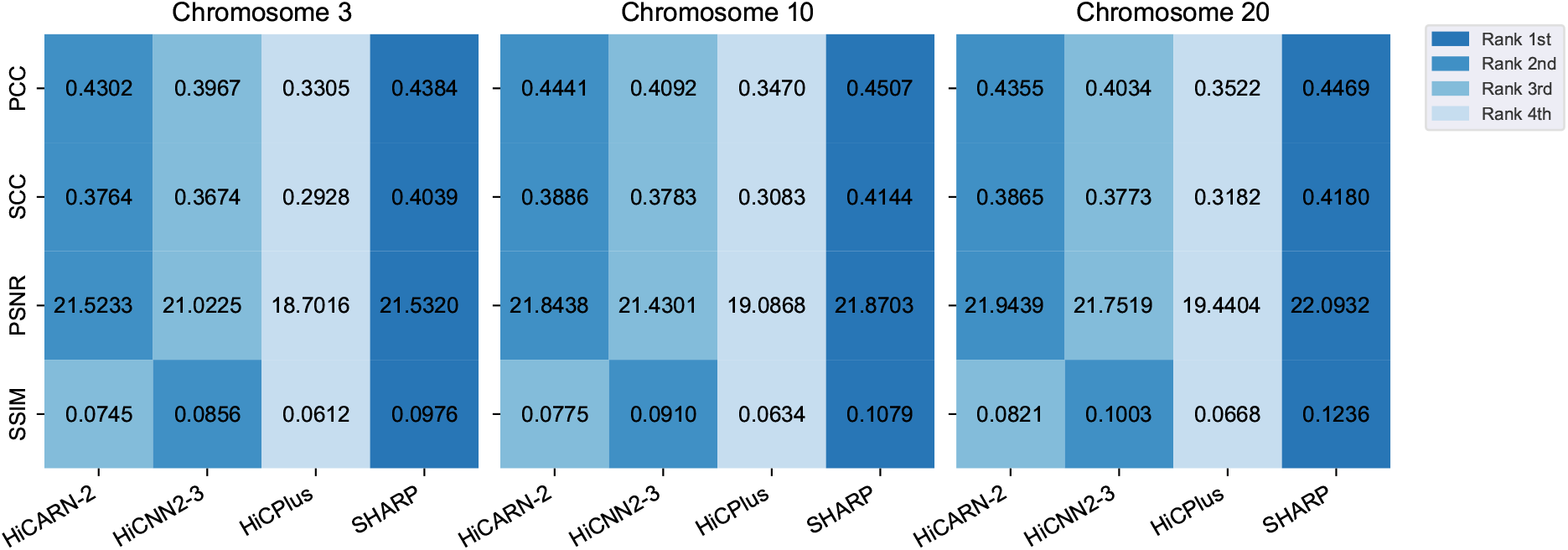
Resolution enhancement performance of SHARP as compared to three other methods when applied to a Hi-C contact matrix of the H1-hESC cell line.

**Supplementary Figure S4:**
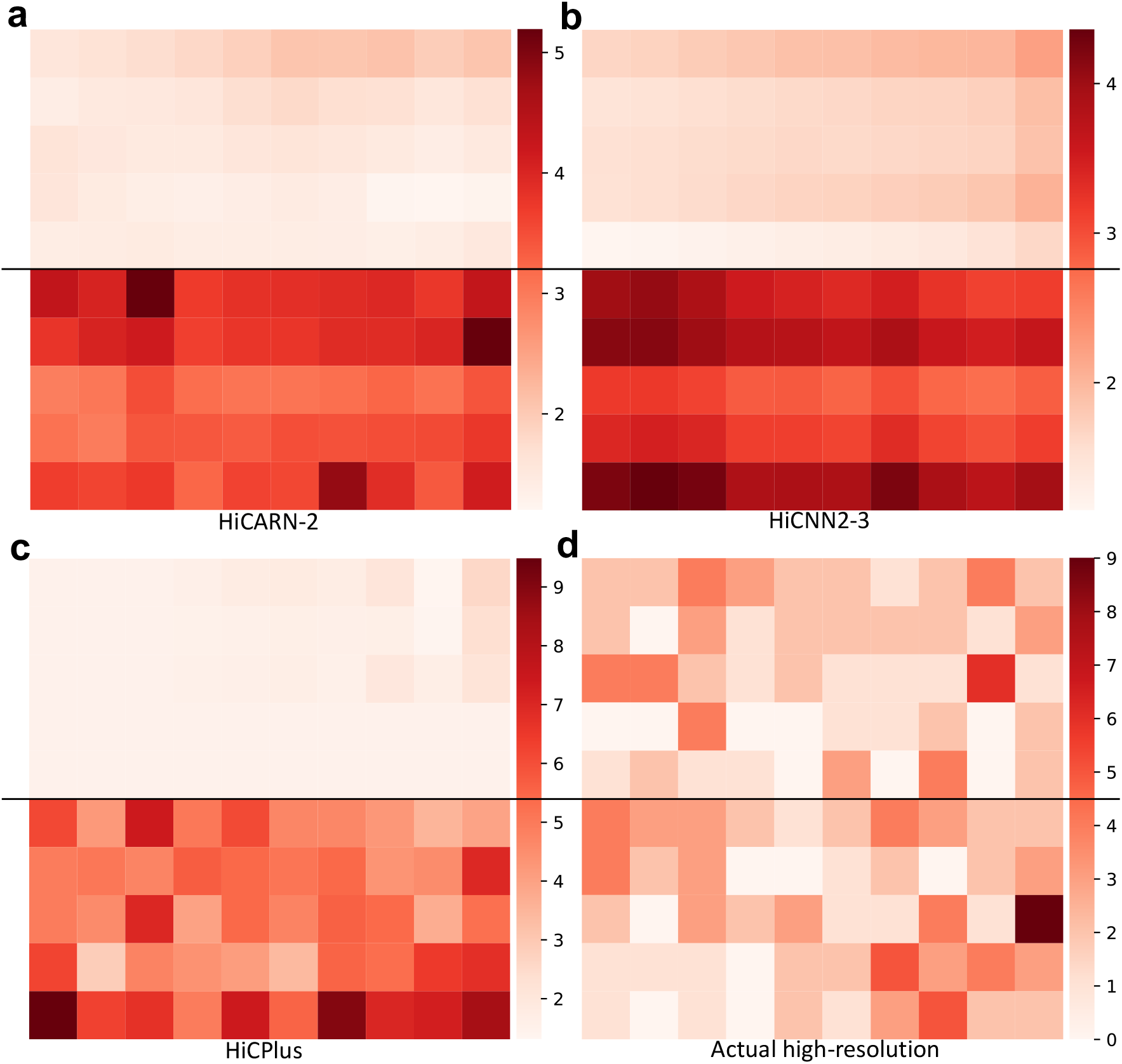
Artificial structures in resolution-enhanced Hi-C contact matrices of chromosome 20 (5.415Mb-5.465Mb *×* 4.045Mb-4.095Mb) of GM12878 cells. **a-c**, The resolution-enhanced contact matrices produced by HiCARN-2 (a), HiCNN2-3 (b), and HiCPlus (c). In all cases, the upper half is assigned to one patch and the lower half is assigned to another patch. **d**, The actual high-resolution contact matrix of the same area.

**Supplementary Figure S5:**
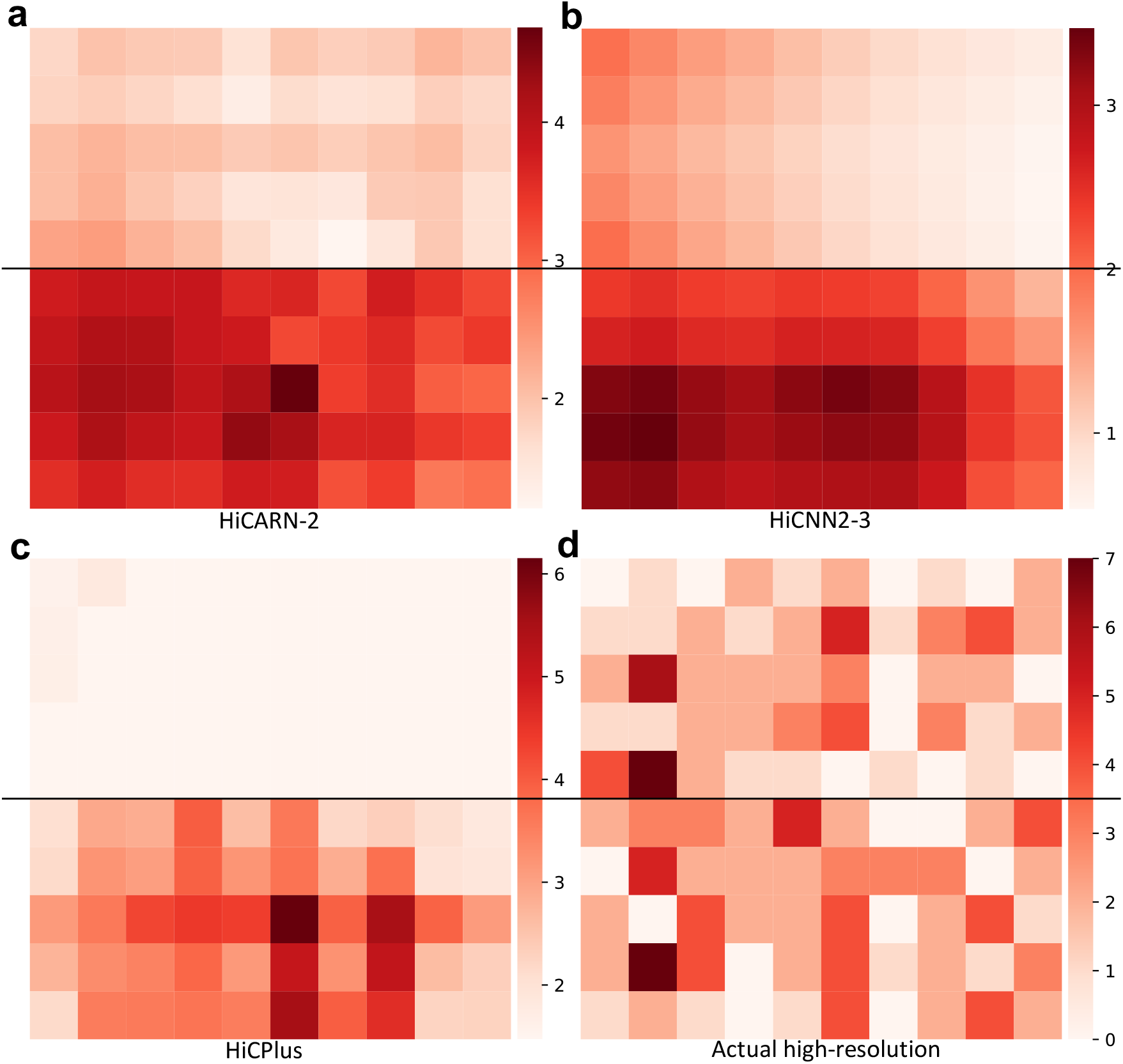
Artificial structures in resolution-enhanced Hi-C contact matrices of chromosome 20 (4.455Mb-4.505Mb *×* 3.46Mb-3.51Mb) of GM12878 cells. **a-c**, The resolution-enhanced contact matrices produced by HiCARN-2 (a), HiCNN2-3 (b), and HiCPlus (c). In all cases, the upper half is assigned to one patch and the lower half is assigned to another patch. **d**, The actual high-resolution contact matrix of the same area.

**Supplementary Figure S6:**
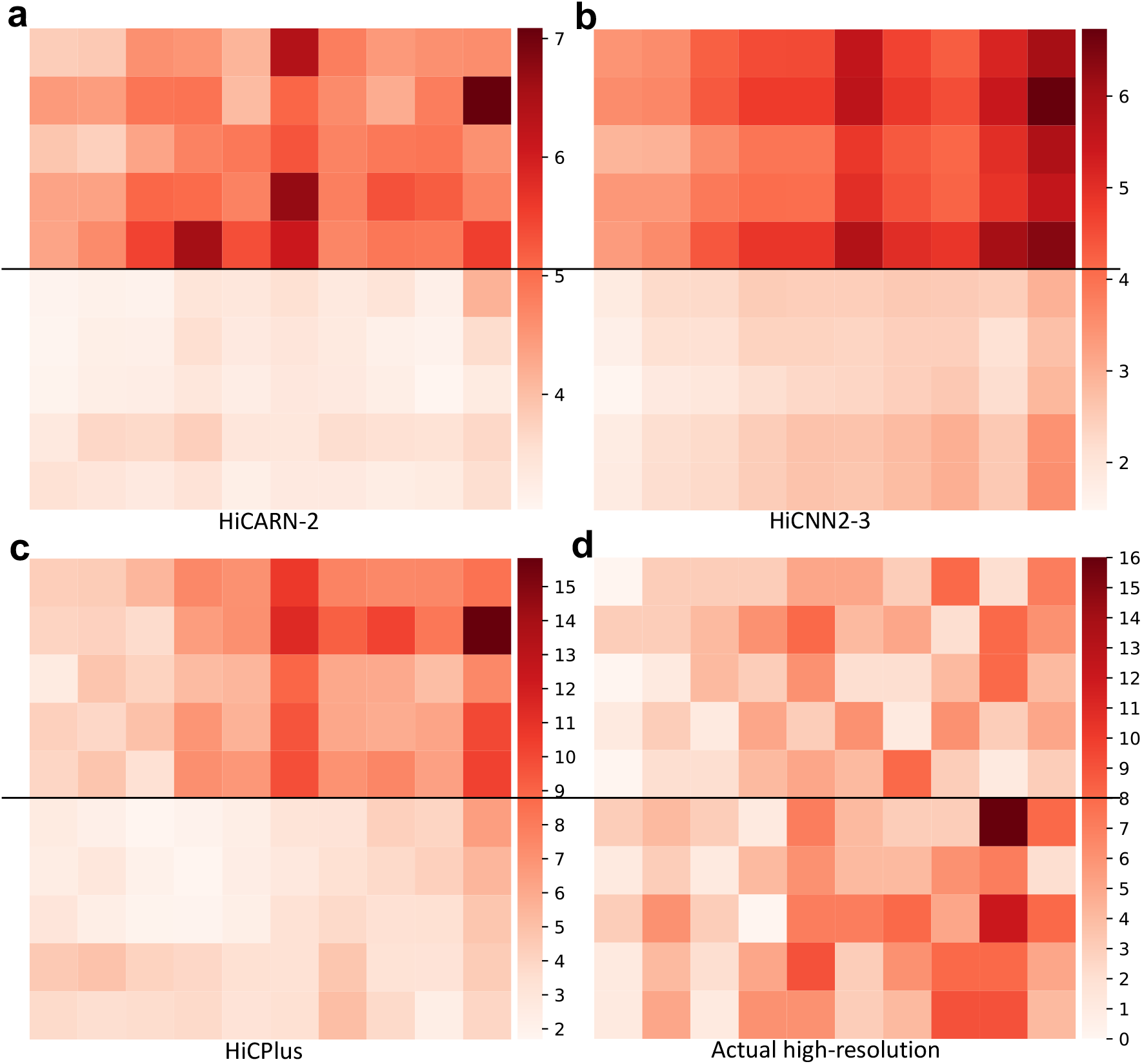
Artificial structures in resolution-enhanced Hi-C contact matrices of chromosome 20 (11.495Mb-11.545Mb *×* 11.85Mb-11.90Mb) of GM12878 cells. **a-c**, The resolution-enhanced contact matrices produced by HiCARN-2 (a), HiCNN2-3 (b), and HiCPlus (c). In all cases, the upper half is assigned to one patch and the lower half is assigned to another patch. **d**, The actual high-resolution contact matrix of the same area.

**Supplementary Figure S7:**
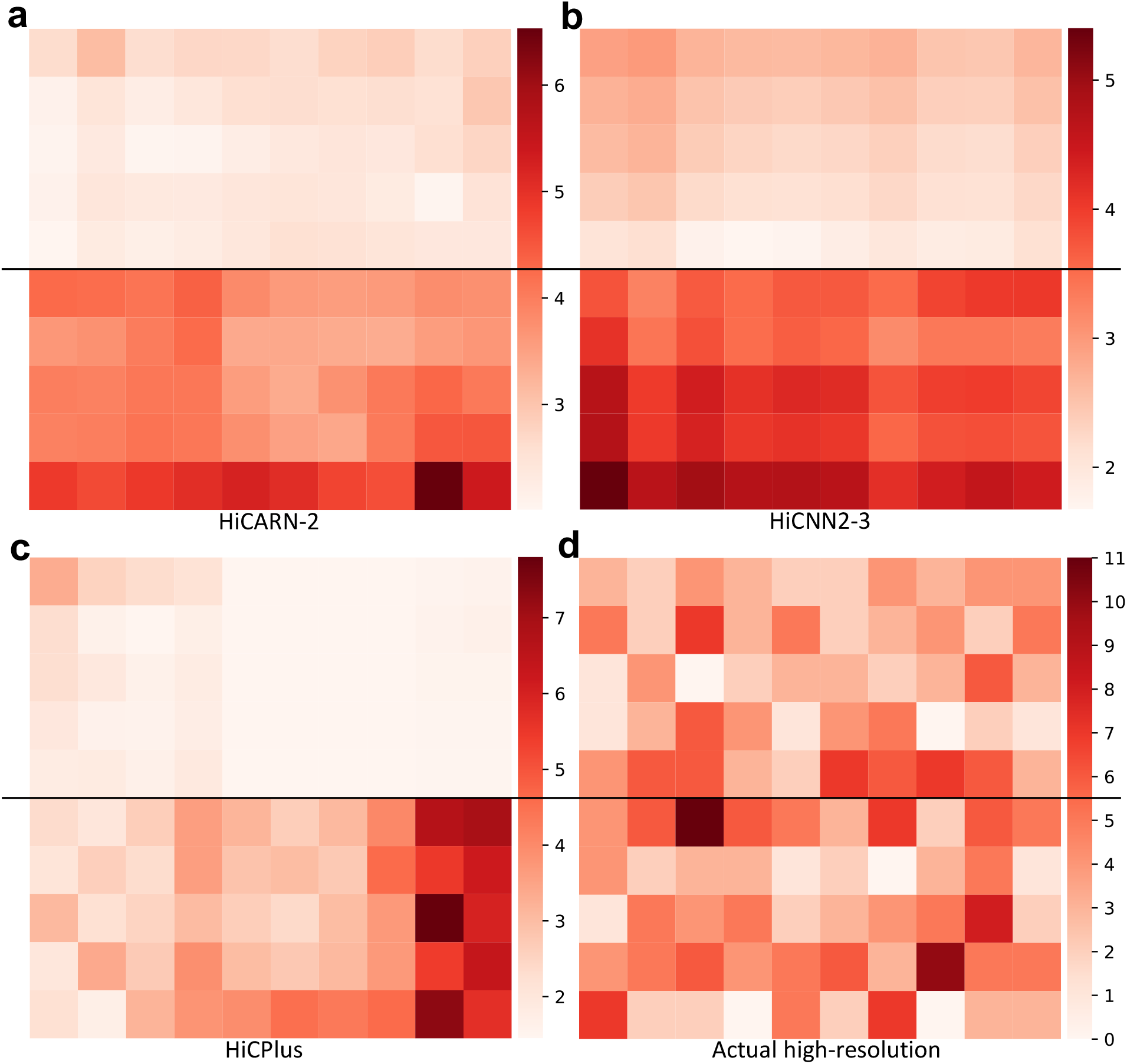
Artificial structures in resolution-enhanced Hi-C contact matrices of chromosome 20 (12.135Mb-12.185Mb *×* 12.69Mb-12.74Mb) of GM12878 cells. **a-c**, The resolution-enhanced contact matrix produced by HiCARN-2 (a), HiCNN2-3 (b), and HiCPlus (c). In all cases, the upper half is assigned to one patch and the lower half is assigned to another patch. **d**, The actual high-resolution contact matrix of the same area.

**Supplementary Figure S8:**
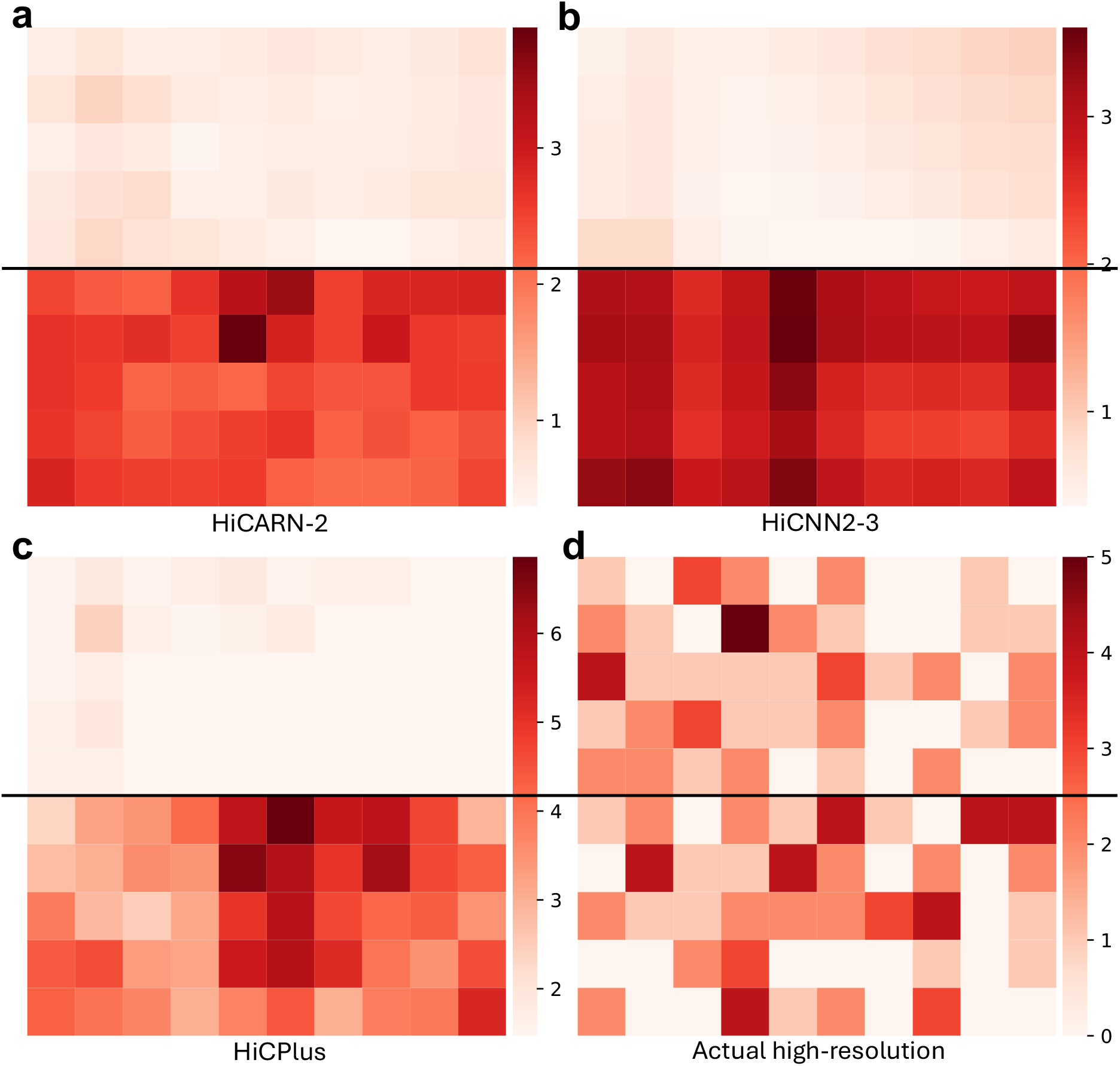
Artificial structures in resolution-enhanced Hi-C contact matrices of chromosome 3 (16.935Mb-16.985Mb *×* 18Mb-18.05Mb) of GM12878 cells. **a-c**, The resolution-enhanced contact matrix produced by HiCARN-2 (a), HiCNN2-3 (b), and HiCPlus (c). In all cases, the upper half is assigned to one patch and the lower half is assigned to another patch. **d**, The actual high-resolution contact matrix of the same area.

**Supplementary Figure S9:**
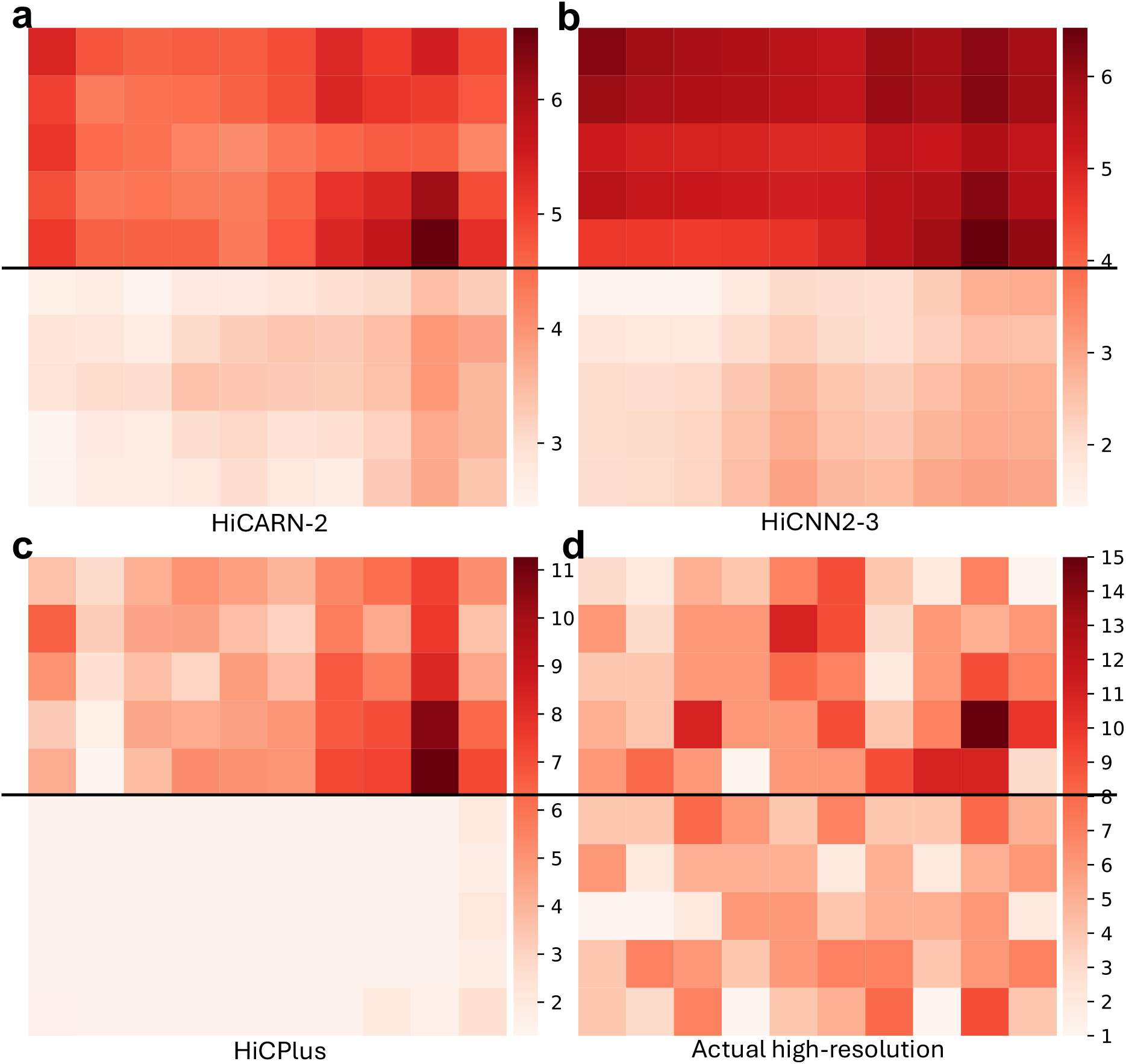
Artificial structures in resolution-enhanced Hi-C contact matrices of chromosome 3 (49.255Mb-49.305Mb *×* 48.51Mb-48.56Mb) of GM12878 cells. **a-c**, The resolution-enhanced contact matrix produced by HiCARN-2 (a), HiCNN2-3 (b), and HiCPlus (c). In all cases, the upper half is assigned to one patch and the lower half is assigned to another patch. **d**, The actual high-resolution contact matrix of the same area.

**Supplementary Figure S10:**
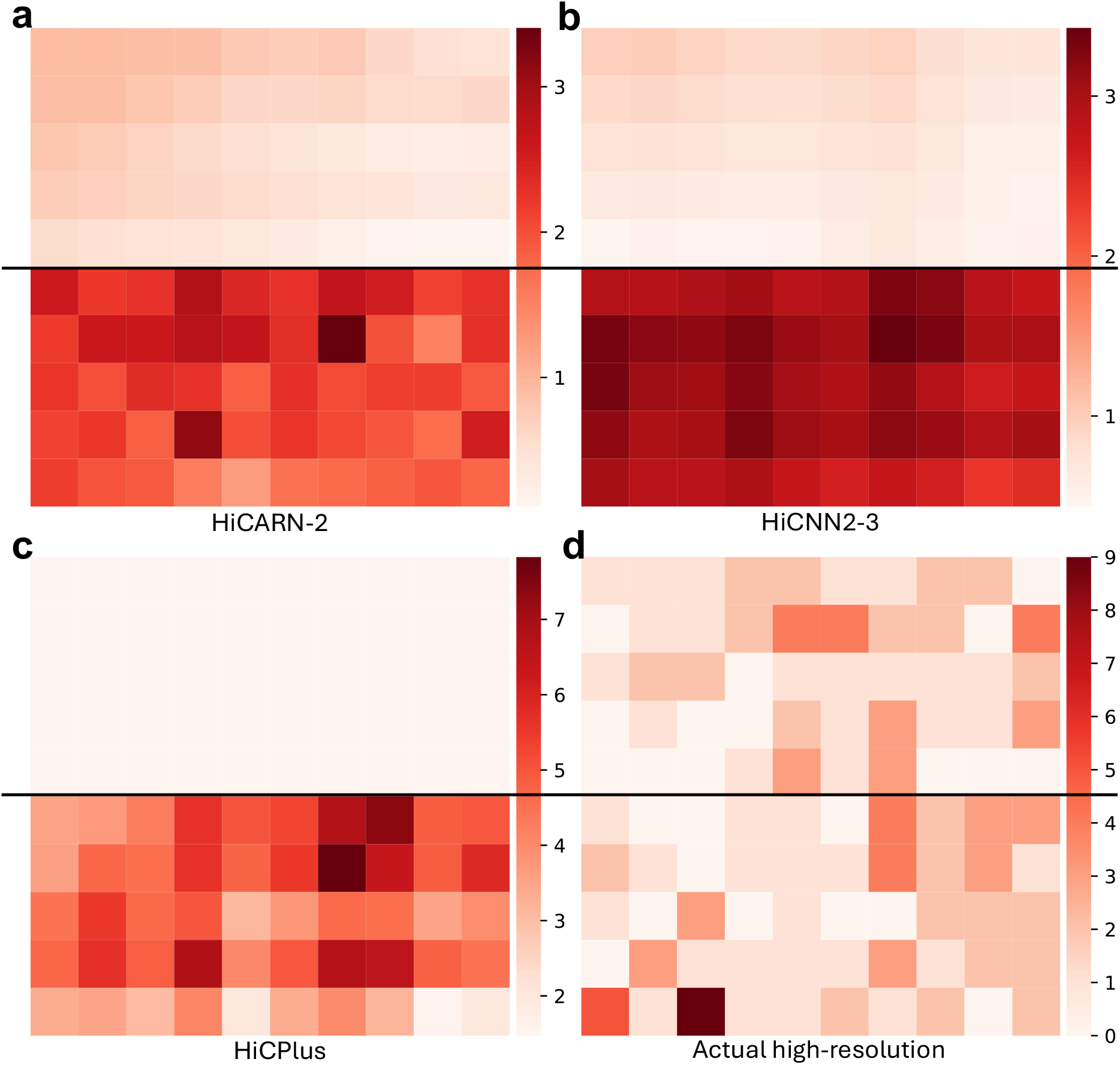
Artificial structures in resolution-enhanced Hi-C contact matrices of chromosome 10 (44.455Mb-44.505Mb *×* 45.255Mb-45.305Mb) of GM12878 cells. **a-c**, The resolution-enhanced contact matrix produced by HiCARN-2 (a), HiCNN2-3 (b), and HiCPlus (c). In all cases, the upper half is assigned to one patch and the lower half is assigned to another patch. **d**, The actual high-resolution contact matrix of the same area.

**Supplementary Figure S11:**
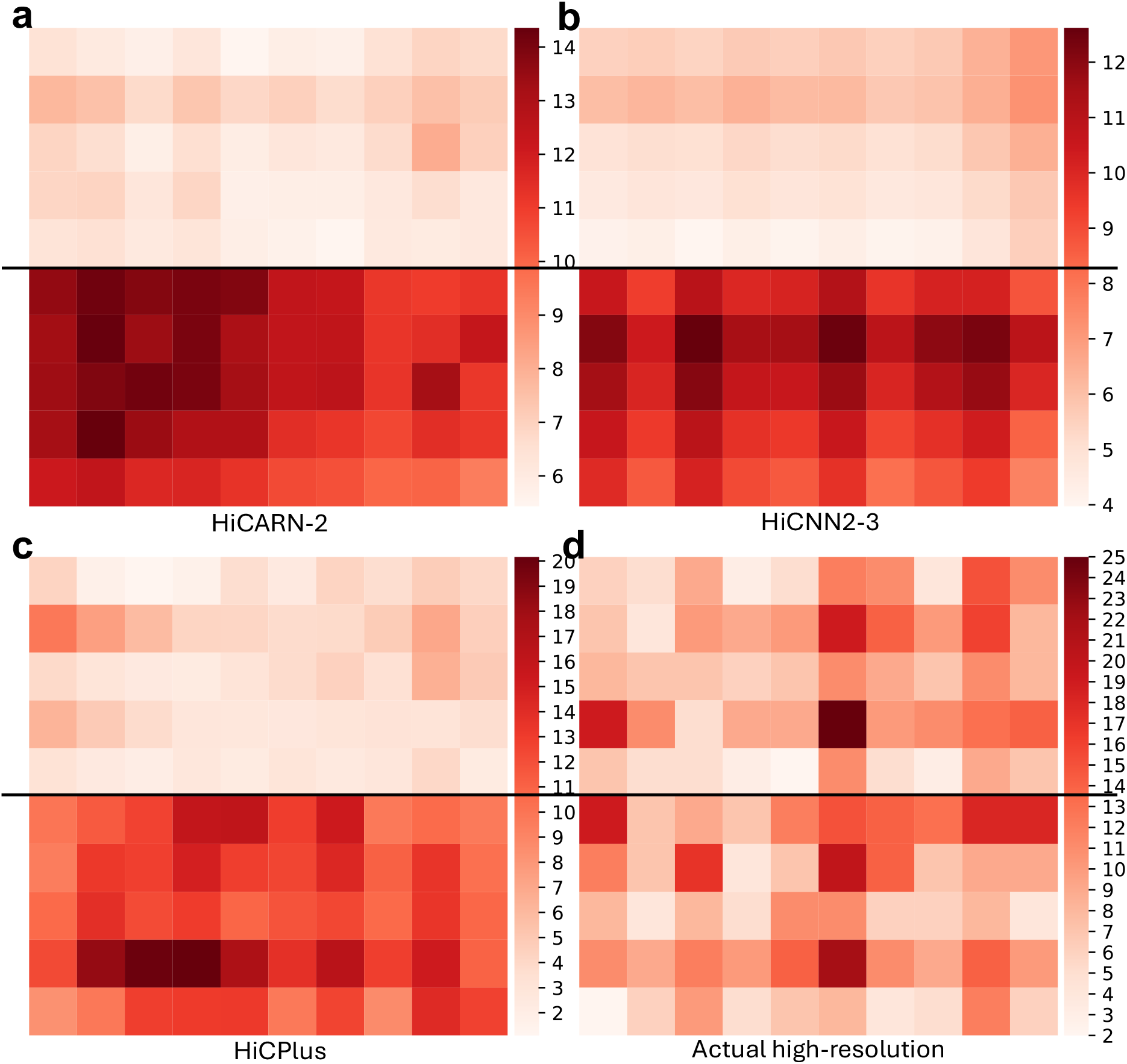
Artificial structures in resolution-enhanced Hi-C contact matrices of chromosome 10 (0.295Mb-0.345Mb *×* 0.665Mb-0.715Mb) of GM12878 cells. **a-c**, The resolution-enhanced contact matrix produced by HiCARN-2 (a), HiCNN2-3 (b), and HiCPlus (c). In all cases, the upper half is assigned to one patch and the lower half is assigned to another patch. **d**, The actual high-resolution contact matrix of the same area.

**Supplementary Figure S12:**
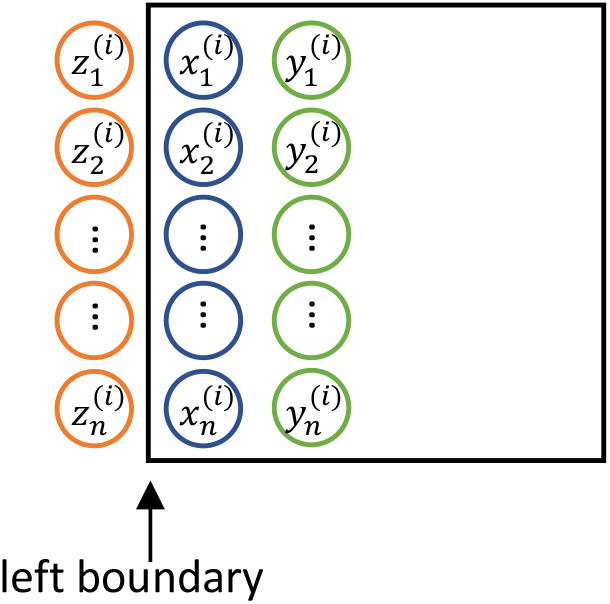
Illustration of the calculation of the NBD measure. The black box shows a patch (Patch *i*) defined by a resolution enhancement method. The NBD measure computes the average difference across a boundary (between *x* and *z*) and uses the average difference of the immediate neighboring values within the patch (between *x* and *y*) as control. The NBD measure considers all four boundaries in this way.

**Supplementary Figure S13:**
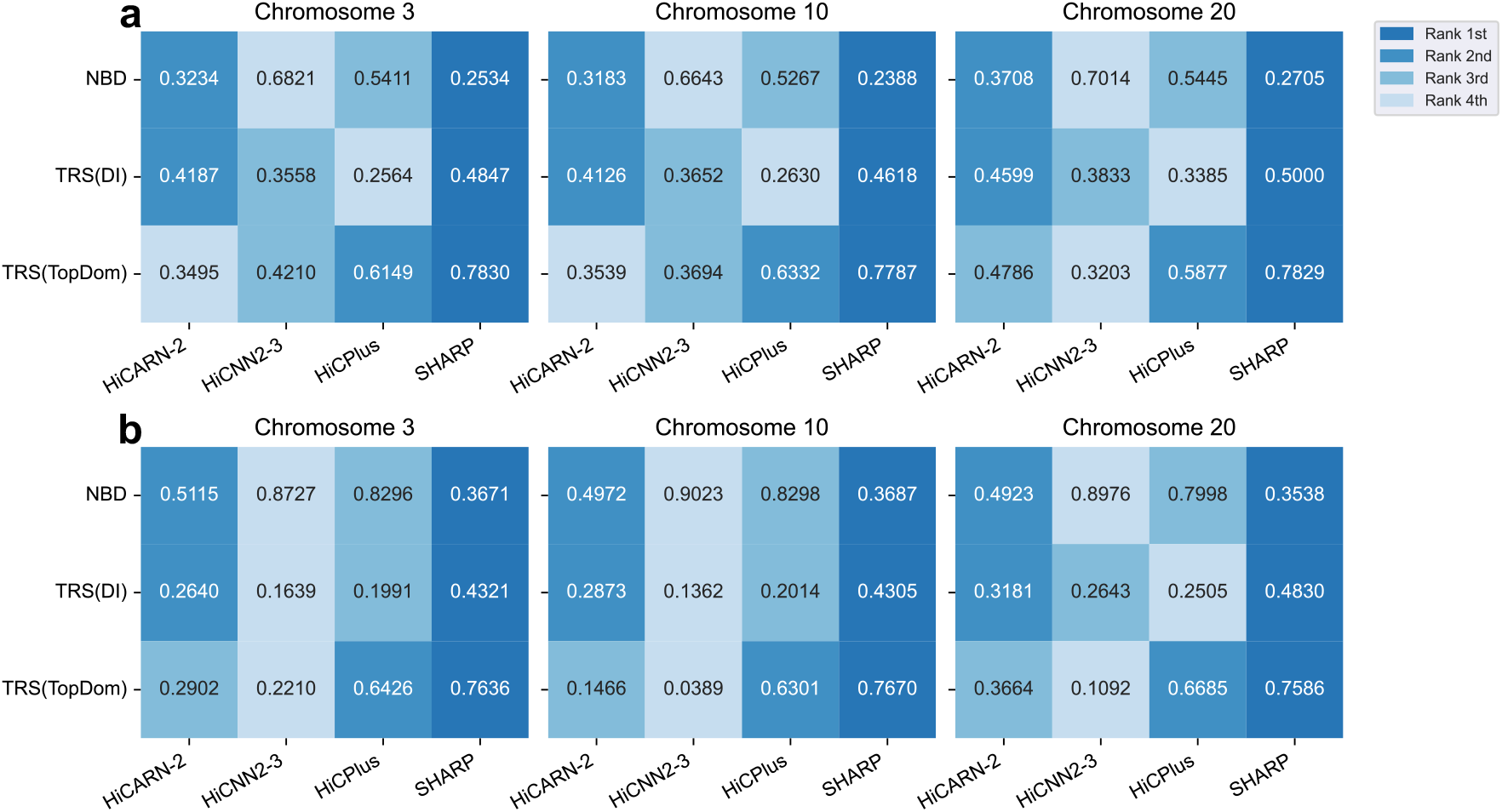
Artificial structure avoidance performance of SHARP as compared to three other methods when applied to low-resolution contact matrices down-sampled from a high-resolution matrix at different sampling rates. For the NBD measure, a smaller value corresponds to better avoidance of artificial structures; for the TRS measures, a larger value corresponds to better avoidance of artificial TADs. **a**, Sampling rate of 1/50. **b**, Sampling rate of 1/100.

**Supplementary Figure S14:**
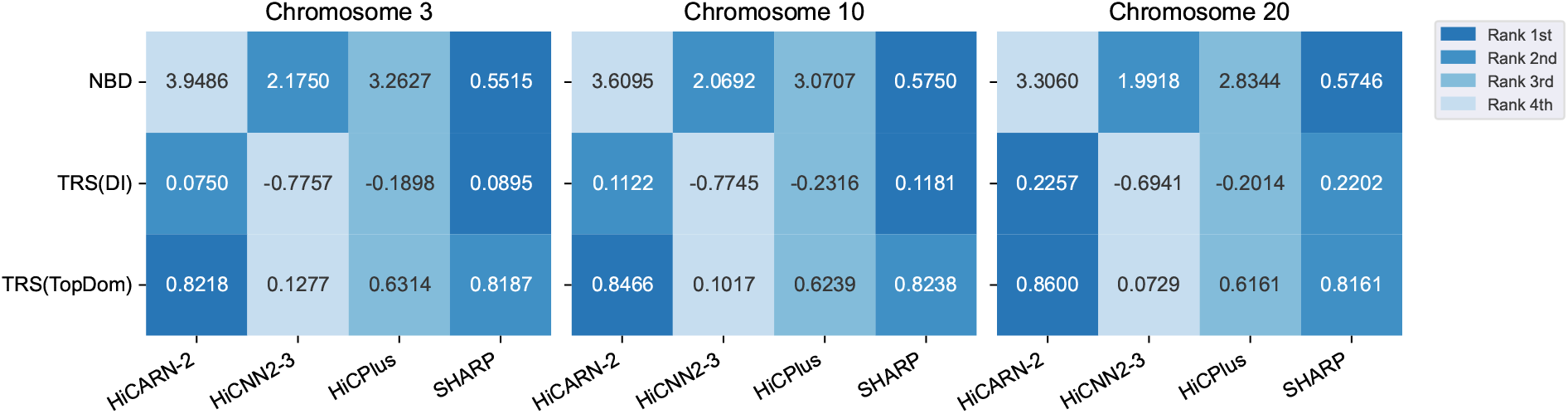
Artificial structure avoidance performance of SHARP as compared to three other methods when applied to an actual low-resolution contact matrix. For the NBD measure, a smaller value corresponds to better avoidance of artificial structures; for the TRS measures, a larger value corresponds to better avoidance of artificial TADs.

**Supplementary Figure S15:**
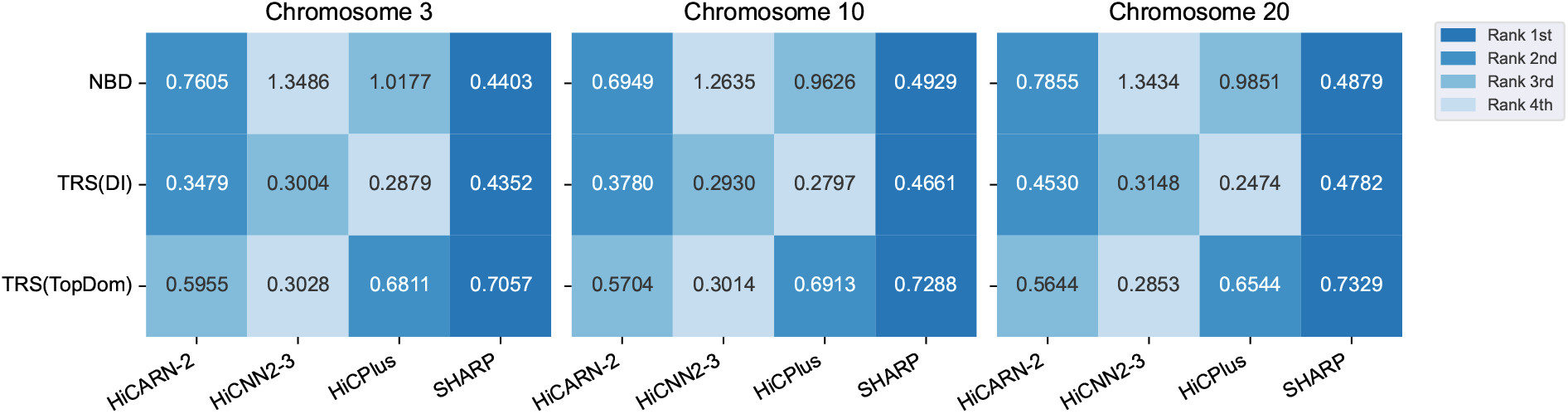
Artificial structure avoidance performance of SHARP as compared to three other methods when applied to a Hi-C contact matrix of the H1-hESC cell line. For the NBD measure, a smaller value corresponds to better avoidance of artificial structures; for the TRS measures, a larger value corresponds to better avoidance of artificial TADs.

**Supplementary Figure S16:**
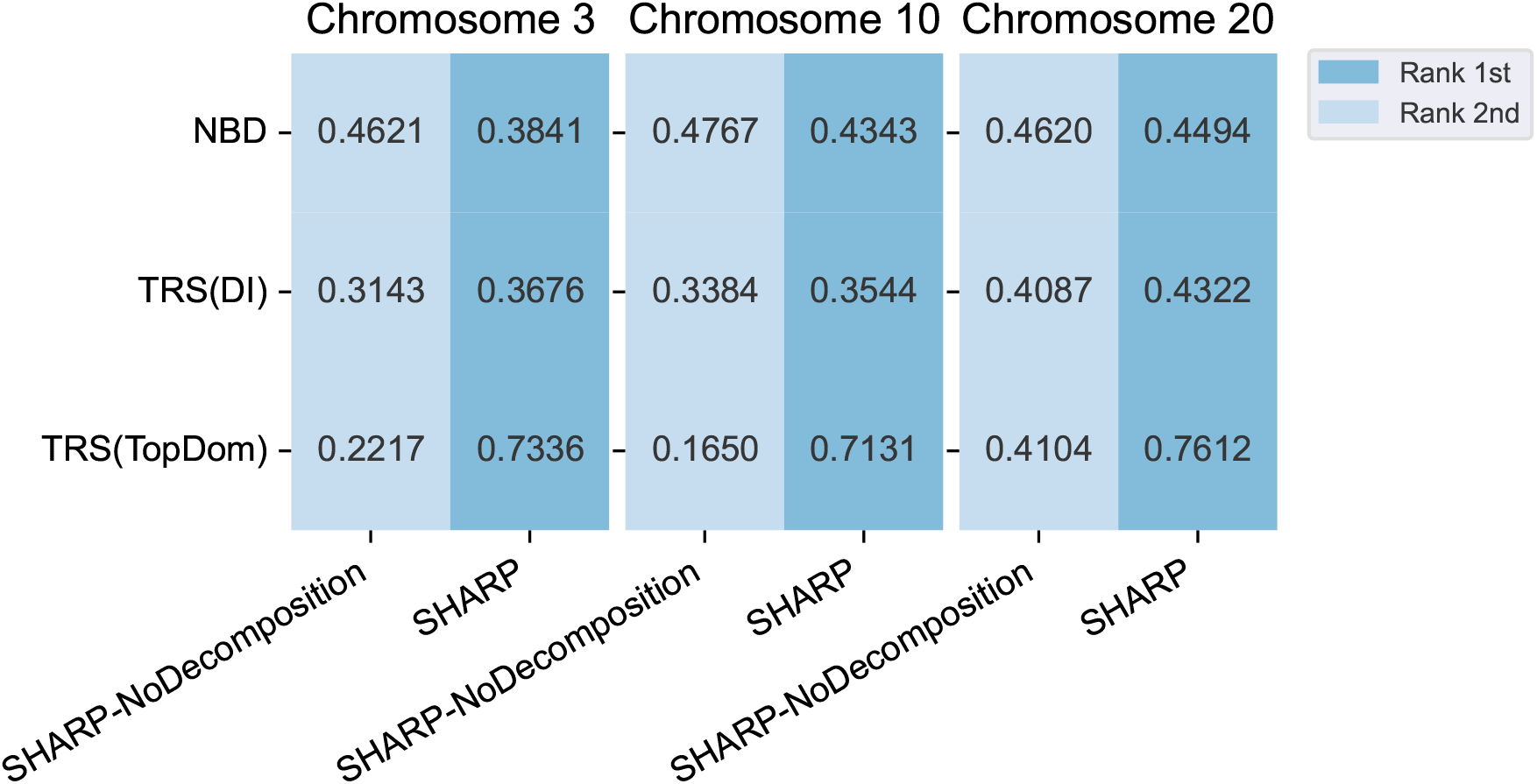
Artificial structure avoidance performance of SHARP and a simplified version of it that does not perform block detection and removal (SHARP-NoDecomposition). For the NBD measure, a smaller value corresponds to better avoidance of artificial structures; for the TRS measures, a larger value corresponds to better avoidance of artificial TADs.

**Supplementary Figure S17:**
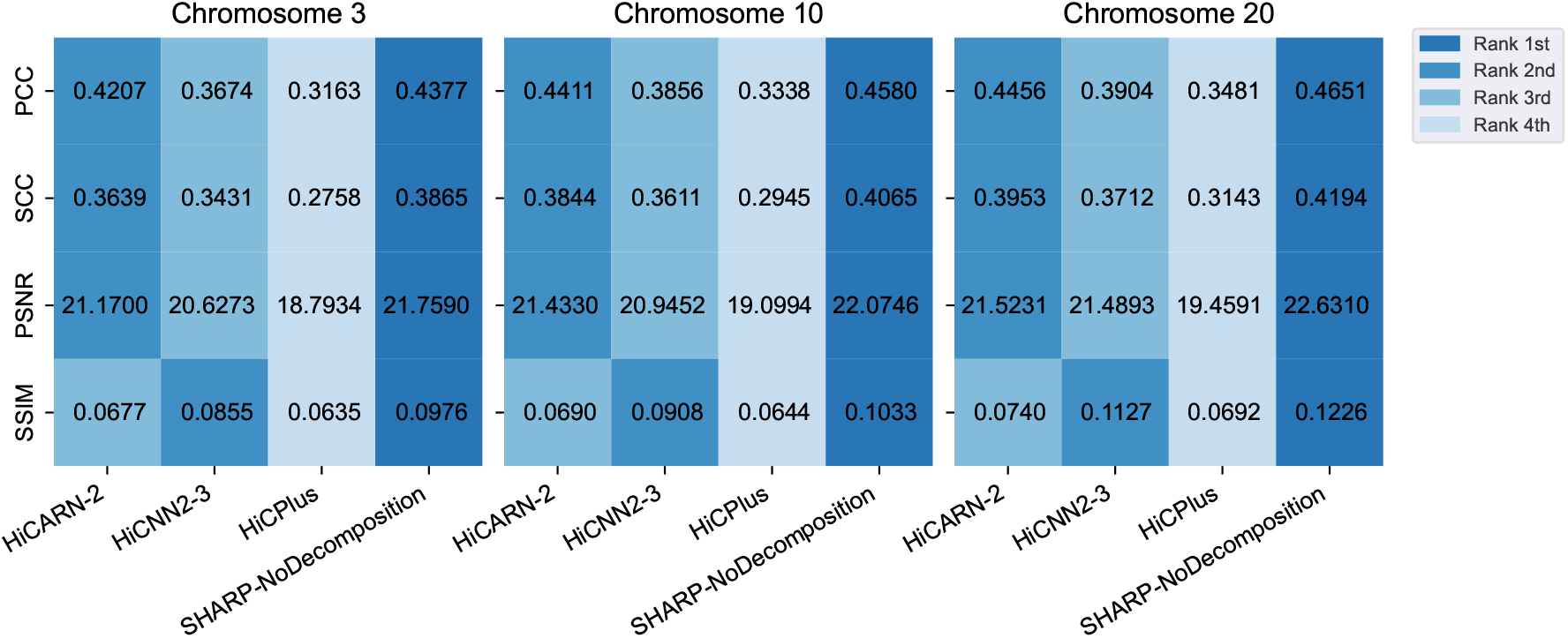
Resolution enhancement performance of SHARP-NoDecomposition as compared to three other methods. For all four measures, a larger value corresponds to better performance.

**Supplementary Figure S18:**
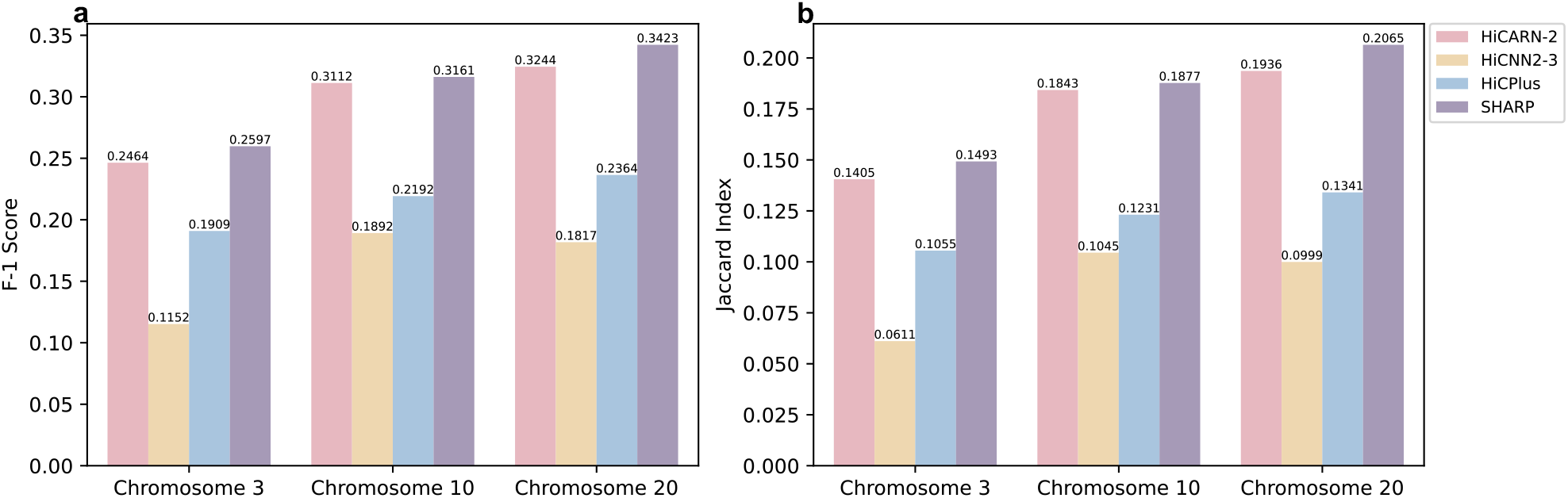
Significant chromatin interactions in the imputed regions (i.e., bin pairs in the evaluation set only) identified by SHARP as compared to three other methods. **a-b**, Comparing the statistically significant chromatin interactions identified from the resolution-enhanced contact matrices produced by the methods with those identified from the actual high-resolution matrix, quantified by the F-1 score (a) or Jaccard Index (b).

**Supplementary Figure S19:**
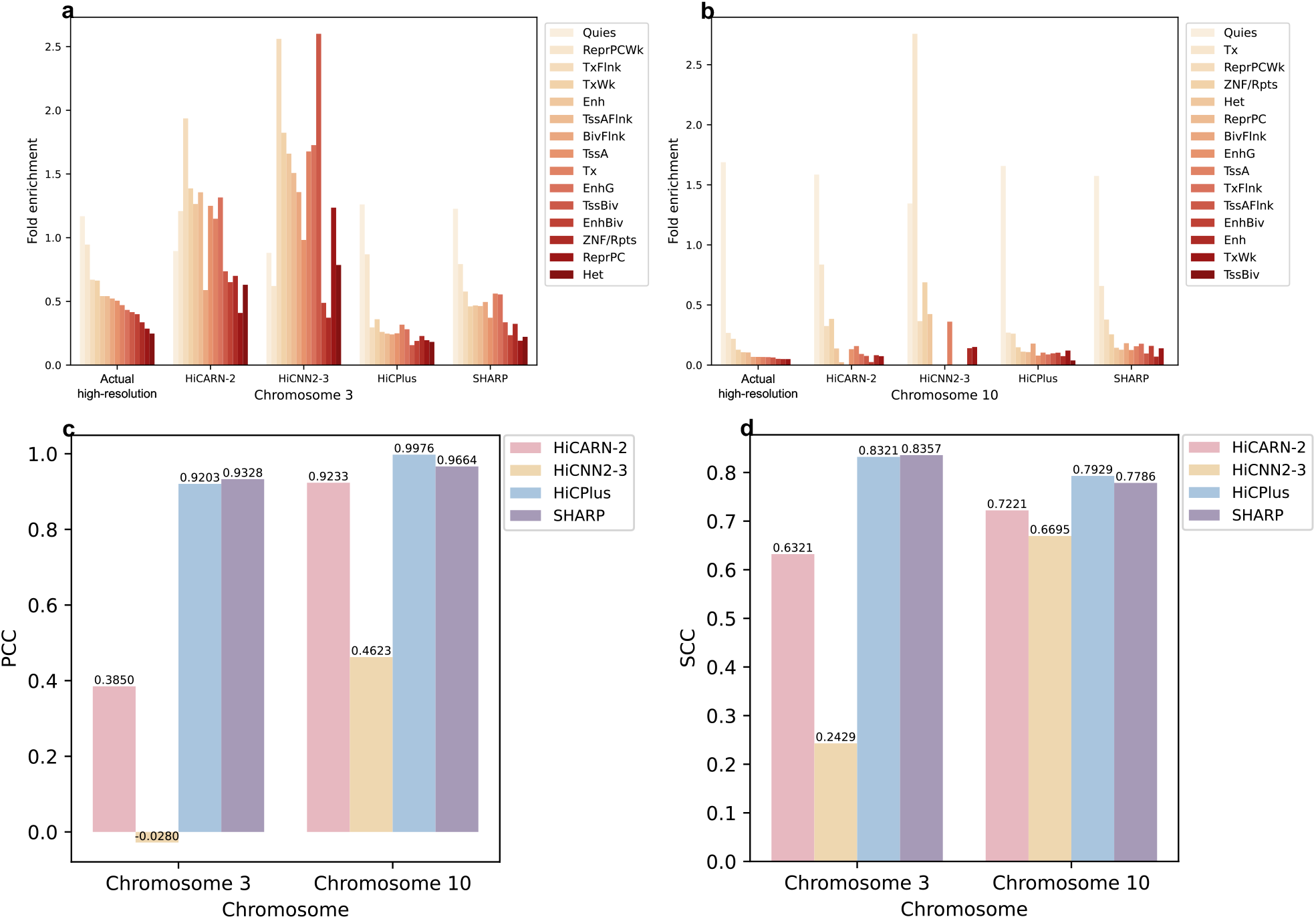
Significant chromatin interactions identified by SHARP as compared to three other methods. **a-b**, Enrichment of the identified chromatin interactions at different chromatin states of Chr 3 (a) and Chr 10 (b). **c-d**, Correlation of the enrichment values with those computed based on significant interactions identified from the actual high-resolution matrix. (c) PCC (d) SCC

**Supplementary Figure S20:**
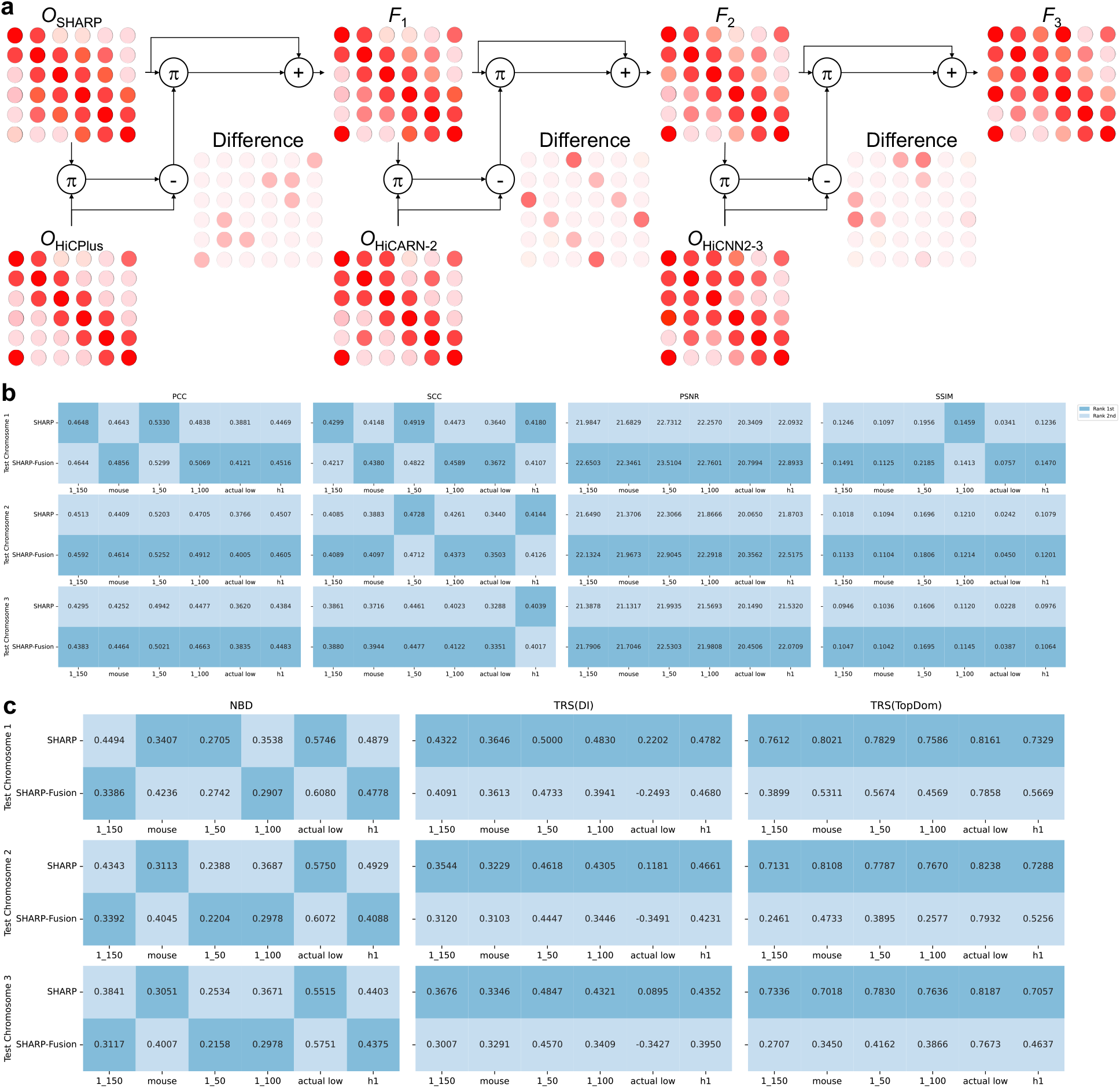
Overview of SHARP-Fusion. **a**, The method of SHARP-Fusion. **b**, Resolution enhancement performance of SHARP-Fusion as compared to SHARP. **c**, Artificial structure avoidance performance of SHARP-Fusion as compared to SHARP.

**Supplementary Figure S21:**
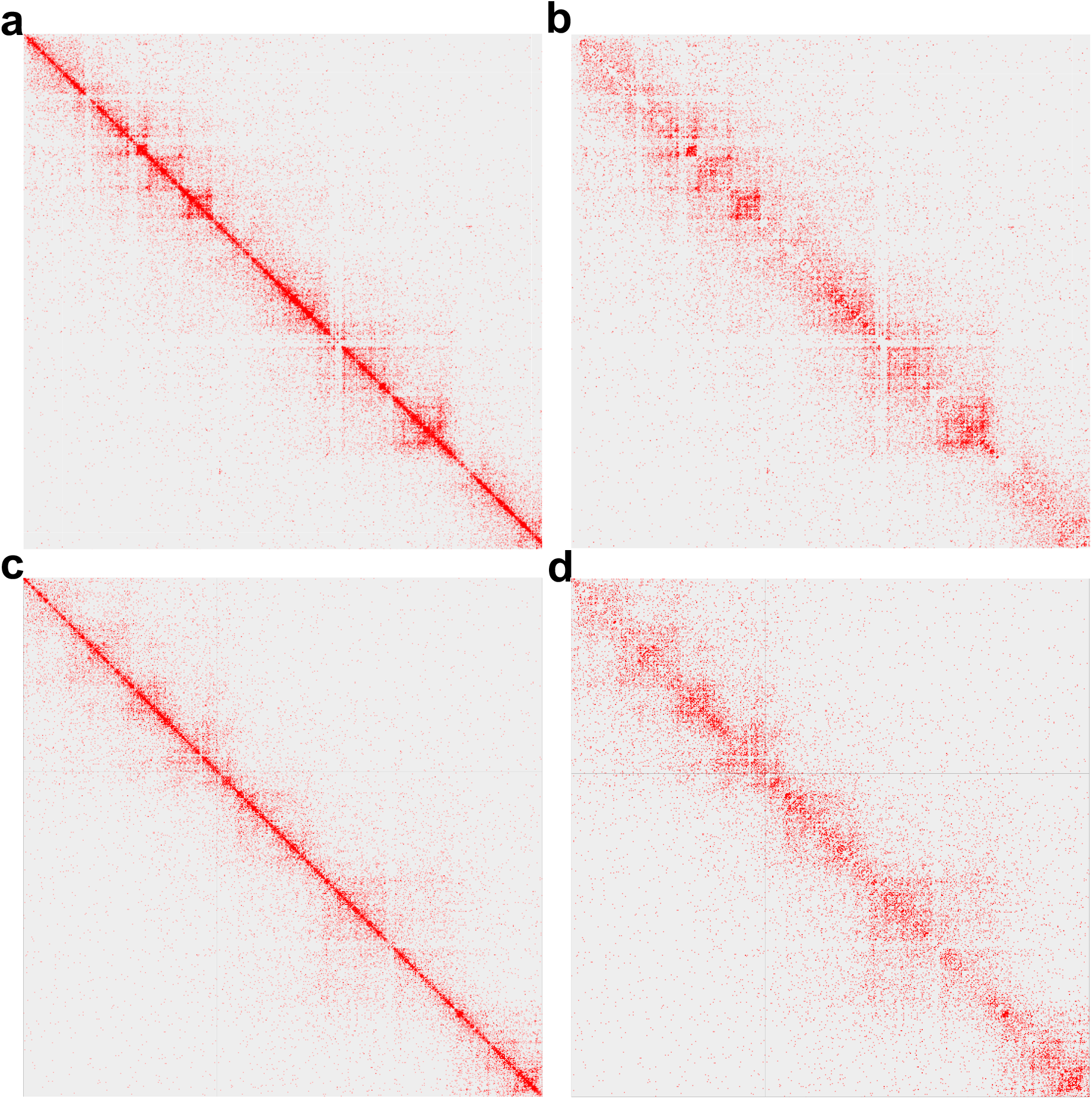
Removing signals due to 1D proximity from the low-resolution GM12878 matrix at subsampling rate of 1/150. **a-b**, The original low-resolution data for the region Chr22:22.3Mb-25Mb (a) and the resulting data after removing signals due to 1D proximity using our formula (b). **c-d**, The original low-resolution data for the region Chr22:36.5Mb-39.2Mb (c) and the resulting data after removing signals due to 1D proximity using our formula (d).

**Supplementary Figure S22:**
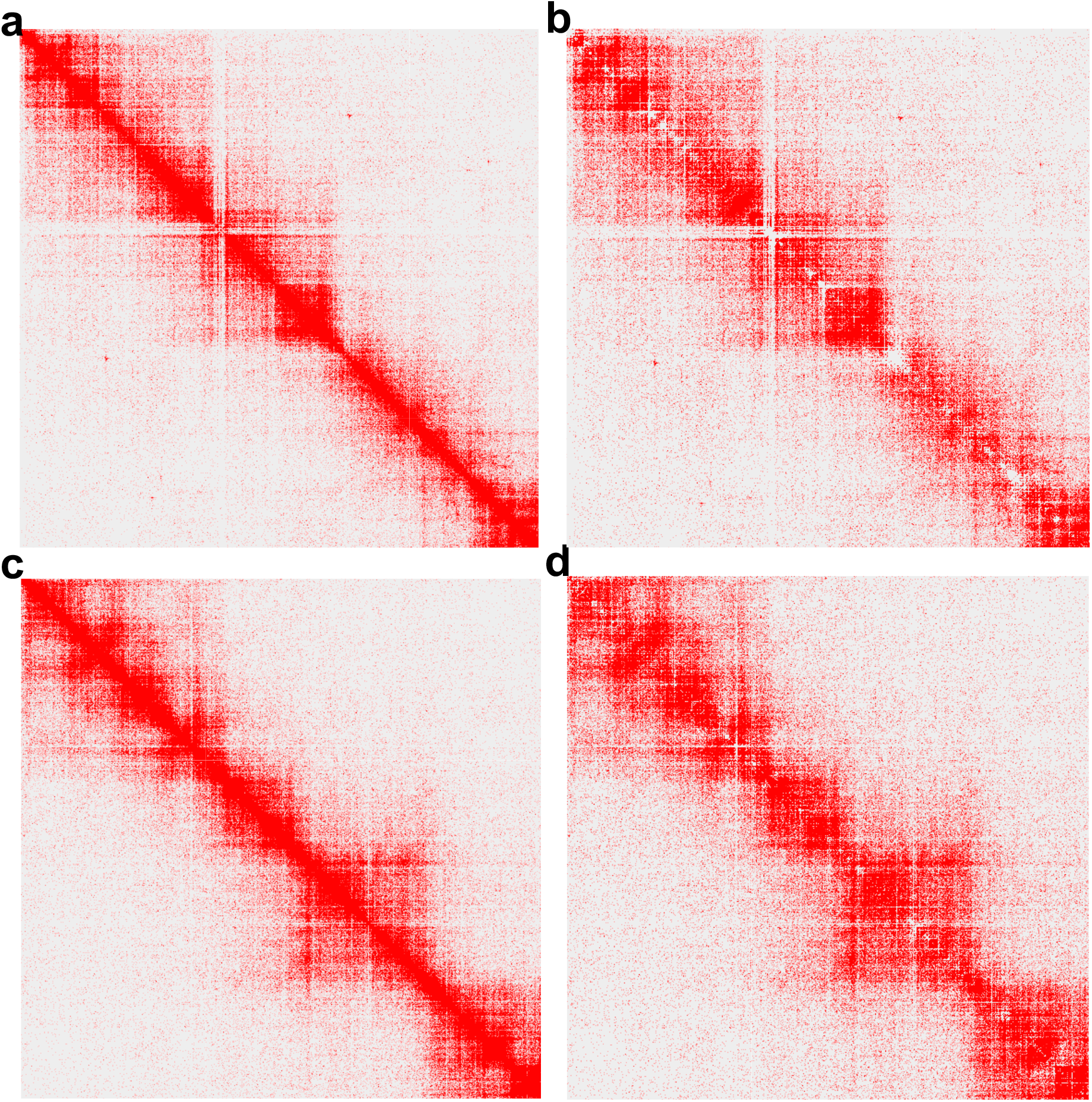
Removing signals due to 1D proximity from the high-resolution GM12878 matrix. **a-b**, The original high-resolution data for the region Chr22:22.3Mb-25Mb (a) and the resulting data after removing signals due to 1D proximity using our formula (b). **c-d**, The original high-resolution data for the region Chr22:36.5Mb-39.2Mb (c) and the resulting data after removing signals due to 1D proximity using our formula (d).

**Supplementary Figure S23:**
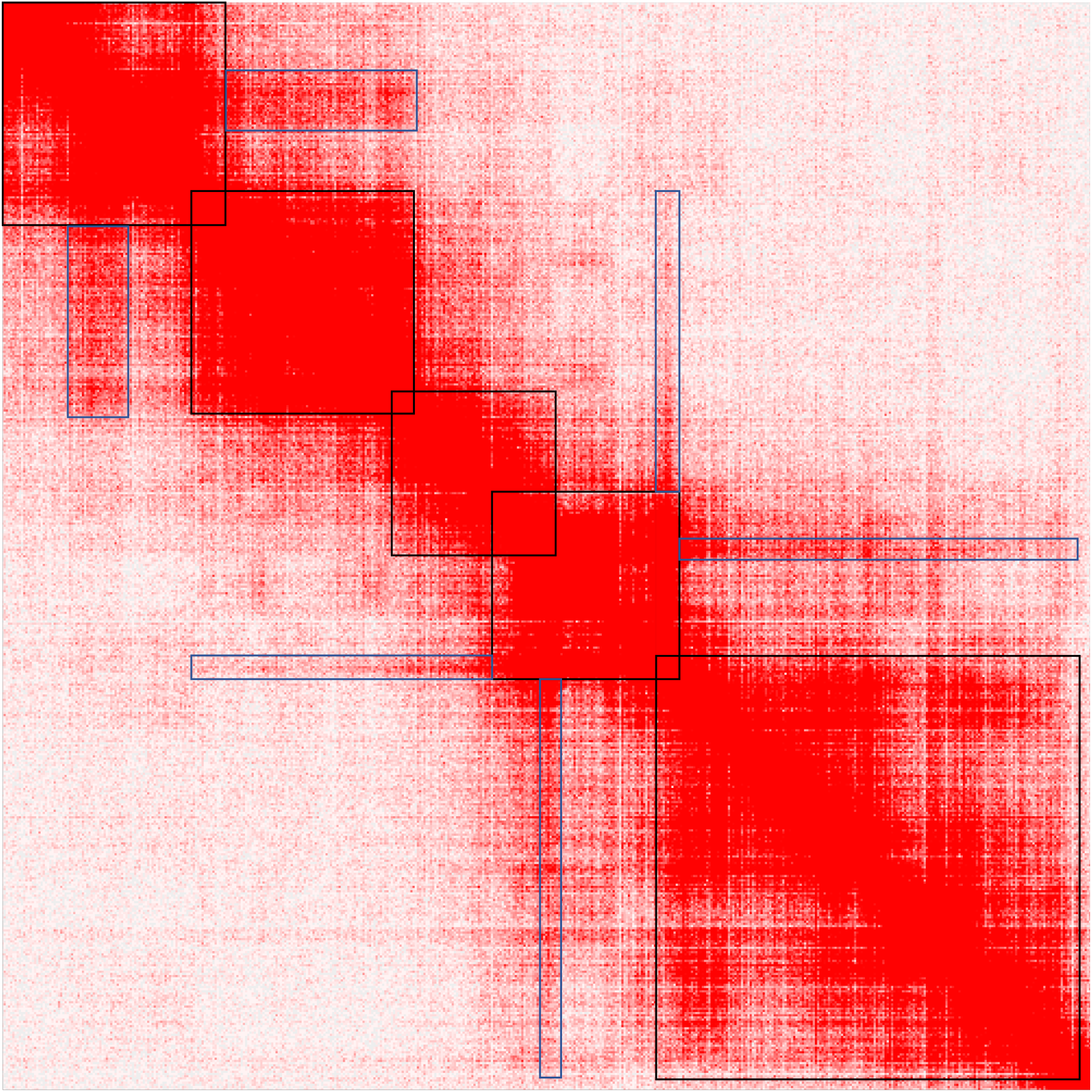
Signals due to contiguous domains. Examples of the within-domain signals on the main diagonal and the between-domain signals off-diagonal are highlighted in black and blue boxes, respectively.

**Supplementary Figure S24:**
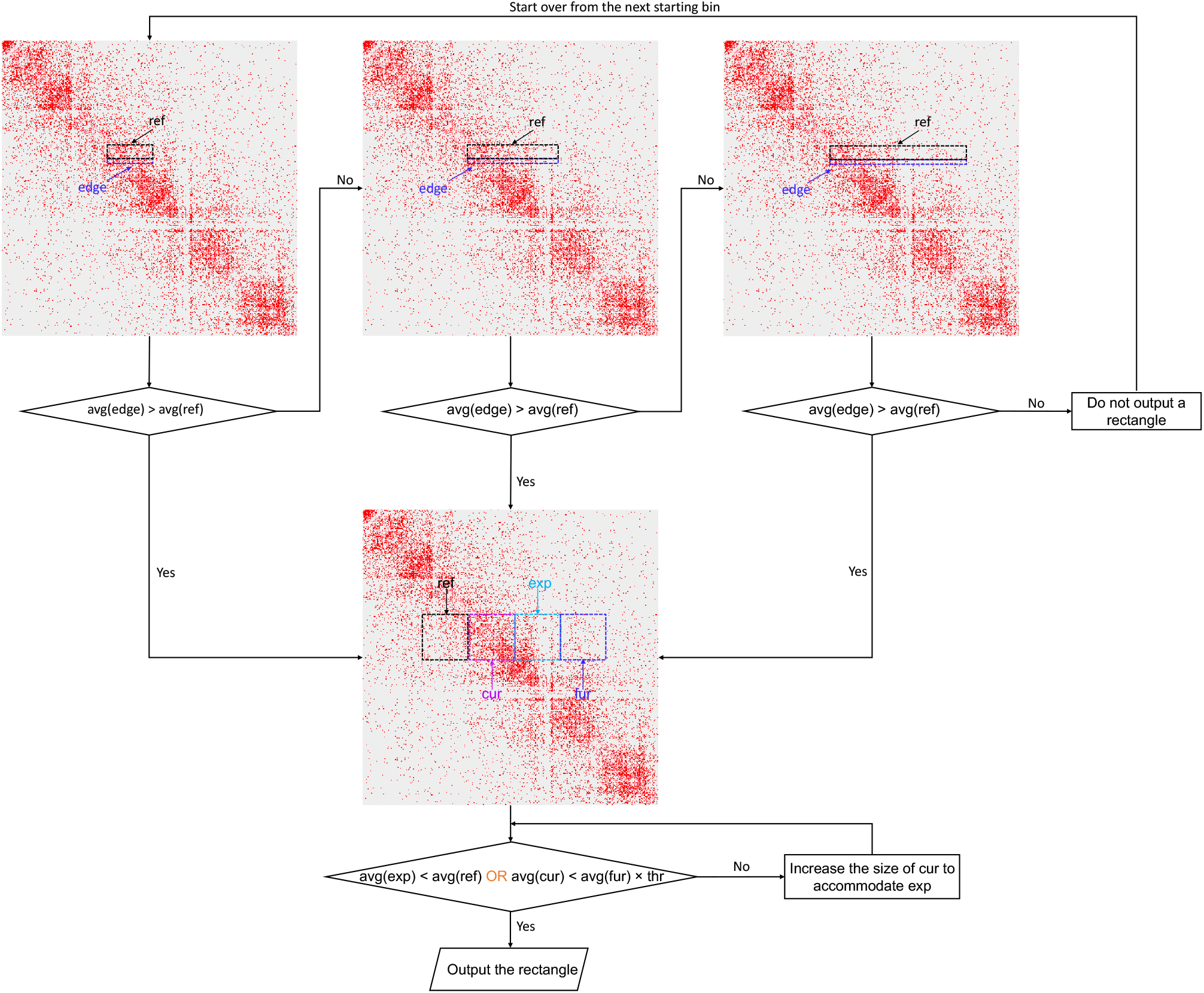
Detection of on-diagonal rectangles caused by contiguous domains. The procedure is illustrated by a flow chart and described in detail in the Methods section.

**Supplementary Figure S25:**
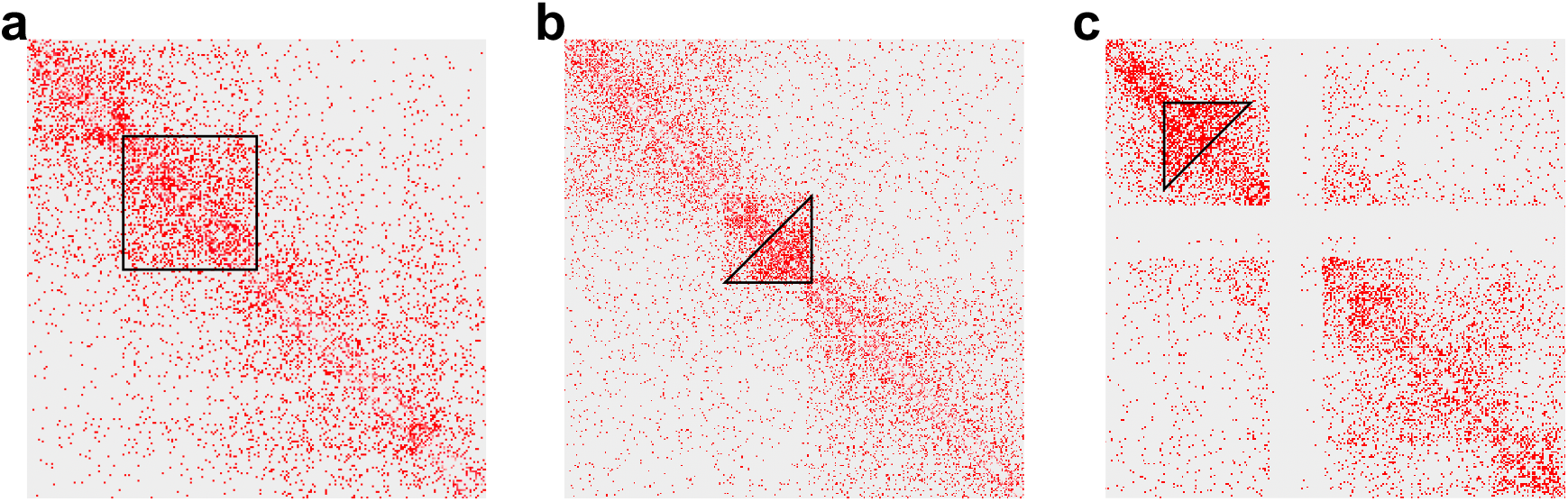
Categorization of on-diagonal rectangles caused by contiguous domains. **a-c**, Common signal patterns of on-diagonal rectangles include uniform signals (a), stronger lower triangle (b), and stronger upper triangle (c).

**Supplementary Figure S26:**
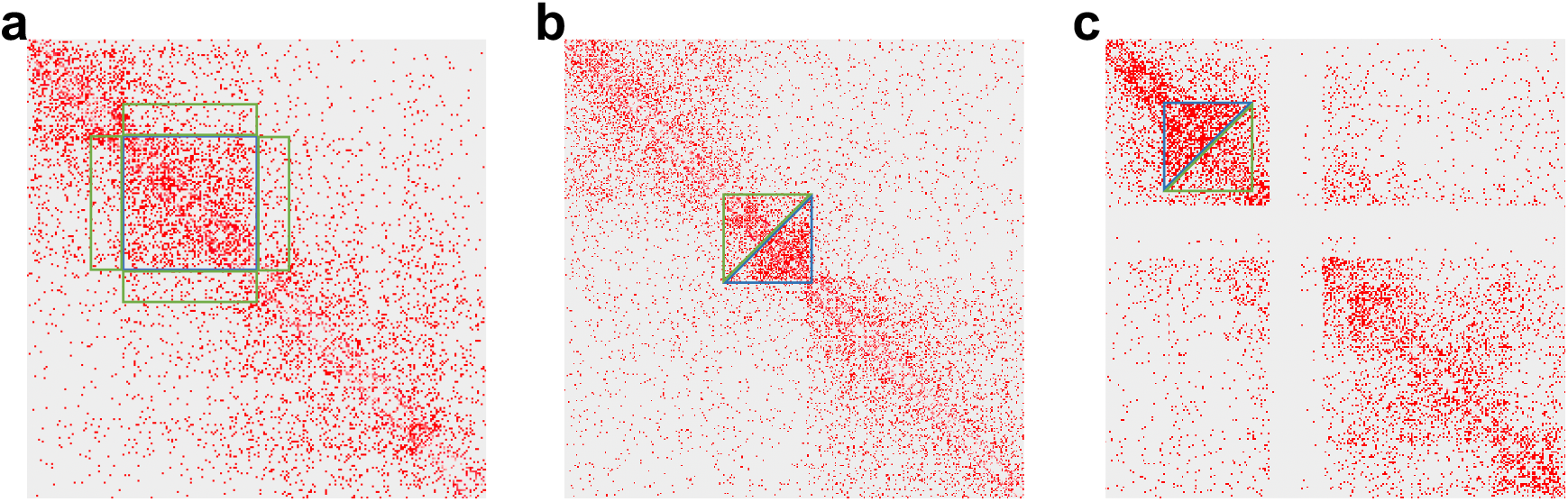
Obtaining strong values and baseline values from on-diagonal blocks. **a-c**, The blue and green lines show areas from which strong values and baseline values are obtained, respectively, for modeling on-diagonal blocks in the uniform signal category (a), lower triangle category (b), and upper triangle category (c).

**Supplementary Figure S27:**
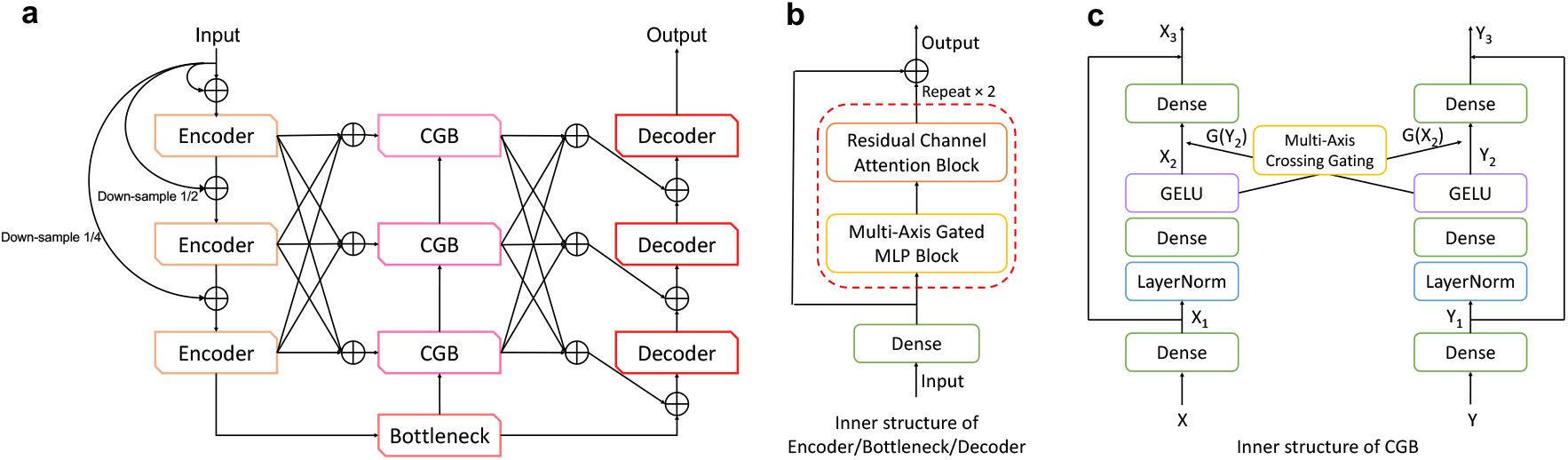
The MAXIM architecture. **a**, The overall design of MAXIM. **b**, Zoom-in view of a encoder, decoder, or bottleneck block. **c**, Zoom-in view of a GCB.

